# An ancient route towards salicylic acid and its implications for the perpetual *Trichormus–Azolla* symbiosis

**DOI:** 10.1101/2021.03.12.435107

**Authors:** Sophie de Vries, Cornelia Herrfurth, Fay-Wei Li, Ivo Feussner, Jan de Vries

## Abstract

Despite its small size, the water fern *Azolla* is a giant among plant symbioses. Within each of its leaflets, a specialized leaf cavity is home to a population of nitrogen-fixing cyanobacteria (cyanobionts). While examples of nitrogen fixing cyanobionts are found across the land plant tree of life, *Azolla* is unique in that its symbiosis is perpetual: the cyanobionts are inherited during sexual and vegetative propagation of the fern. What underpins the communication between the two partners? In angiosperms, the phytohormone salicylic acid (SA) is a well-known regulator of plant–microbe interactions. Using HPLC-MS/MS, we pinpoint the presence of SA in the fern; using comparative genomics and phylogenetics, we mined homologs of SA biosynthesis genes across Chloroplastida (Viridiplantae). While canonical isochorismate synthase (ICS) sequences are largely limited to angiosperms, homologs for the entire Phenylalanine ammonia-lyase (PAL)-dependent pathway likely existed in the last common ancestor of land plants. Indeed, *A. filiculoides* secondarily lost its ICS, but has the genetic competence to derive SA from benzoic acid. Global gene expression data from cyanobiont-containing and -free *A. filiculoides* unveil a putative feedback loop: SA appears to induce cyanobacterial proliferation, which in turn down-regulates genes in SA biosynthesis and its responses.

## INTRODUCTION

The water fern *Azolla filiculoides* is best-known for its unique symbiosis with a nitrogen-fixing cyanobacteria, hereafter referred to as the cyanobionts (Rai et al., 2000). Unlike any other nitrogen-fixing symbiosis, the one of *Azolla* and its cyanobiont is vertically inherited from generation to generation (Peters and Meeks 1989, Zheng et al. 2009a, de Vries and de Vries 2018). Indeed, the cyanobiont cannot live without its host (Peters and Meeks 1989, Zheng et al. 2009b). This dependency is written in its genome, too (Ran et al. 2010): the genome has undergone a strong reduction and erosion. On the other side, when well-supplied with nitrogen nutrients *Azolla* is—in the lab—capable of living without its cyanobionts (Shi and Hall 1988). Yet, in nature the two partners occur together.

The *Azolla* cyanobiont is a section IV (Rippka et al. 1979), filamentous nitrogen-fixing cyanobacterium; taxonomically it is most commonly assigned to the genus *Anabaena* or *Nostoc*—but this affiliation is still debated (Pereira & Vasconcelos 2014; de Vries and de Vries 2018). As of late it is called *Trichormus azollae* (Pereira & Vasconcelos 2014). The cyanobiont lives in the leaf cavities of *Azolla* species (Peters and Meeks 1989) and co-occurs with several other bacterial associates, some which are denitrifiers, likely benefitting from the cyanobiont (Dijkhuizen et al. 2018). Additionally, some of the co-occurring bacteria appear to be inherited together with the cyanobiont (Zheng et al. 2009b).

The life cycle of *Azolla* is intertwined with that of its cyanobiont (Rai et al. 2000; Zheng et al. 2009a; de Vries and de Vries, 2018). During vegetative propagation (i.e. simple growth of the fern body and break-up into clonal “colonies”), cyanobiont populations are transferred to new leaf cavities via specialized trichomes (Calvert et al. 1985; Hill 1989). The trichomes grow out of the newly forming leaf cavities and get associated to the shoot apical meristem (Calvert and Peters 1981, Zheng et al. 2009b). On the apical meristem is a colony of hormogonia-like filaments of the cyanobiont (Peters et al. 1978, Peters and Meeks 1989); these filaments become attached to the branched trichome and as the leaf grows, both the trichome and the cyanobiont filaments are engulfed by the formed leaf cavity (Peters and Meeks 1989, Zheng et al. 2009a). Whether the other bacteria are transferred by the same means and whether as a result each leaf cavity is equipped with a highly similar microbiome is currently not known. During sexual propagation, the cyanobionts are attracted to the developing sporocarps, where they, similar to the vegetative propagation, become entangled in trichomes (Becking, 1987; Perkins and Peters 1993). While the formation of the indusium cell layer around the spores occurs, the cyanobionts are moved upwards to the tips of the sporocarps, where they enter through the indusium pore into the forming indusium chamber (Perkins and Peters 1993). Upon maturation of the sporocarp, the cyanobionts in this chamber turn into resilient akinetes, facilitating the co-dormancy of symbiont and host (Perkins and Peters 1993, Zheng et al. 2009b). Hence, the process of sporocarp maturation and cyanobiont differentiation appears to be tightly coordinated (Zheng et al. 2009b). The molecular mechanisms that underpin this coordination are however unknown.

Vegetative propagation has been studied with regard to its chemical set-up. It appears that especially the trichomes are rich in phenolic compounds (Pereira & Carrapiço 2007). And it has been hypothesized that these phenolics may be relevant in some way for the transfer of cyanobionts from one cavity to another (Pereira & Carrapiço 2007). Indeed, it has been highlighted that the expression of a *chalcone synthase* (*CHS*) homolog of *A. filiculoides* is both responding to the absence of nitrogen and the absence of cyanobionts from the leaf pockets (Li et al. 2018, Eily et al. 2019). A recent study identified homologous genes encoding enzymes of the flavonoid biosynthesis pathway and functionally characterized a key enzyme, leucoanthocyanidin reductase (LAR) (Güngör et al. 2021). The identified homologs of the flavonoid biosynthetic genes (homologs of *chalcone isomerase* (*CHI*), *CHS* and *dihydroflavonol 4-reductase* (*DFR*) and *AfLAR*) appear co-expressed during the diurnal cycle of *A. filiculoides* independent of the nitrogen supply (Güngör et al. 2021). However, without nitrogen the genes rapidly decrease in expression level during the night cycle (Güngör et al. 2021).

Among the flavonoids that have been identified in *A. filiculoides* and also another water fern, *Azolla pinnata*, are derivatives of Caffeoylquinic acid, Dicaffeoylquinic (Epi-)Catechin, Quercitin and Kaempferol (Güngör et al. 2021). These flavonoids are involved in defense responses in other plants, e.g. the gymnosperms *Picea abies* (Danielsson et al. 2011). Similarly, another phenolic compound involved in plant defense, salicylic acid (SA) (Ding and Ding 2020), has also been implicated in altering the abundance of the cyanobionts and its expression of genes involved in nitrogen fixation (de Vries et al. 2018). This may suggest that a link between this highly coordinated symbiosis and the defense capabilities of *Azolla* has evolved in the 66-100mya old association (Hall and Swanson 1968, Collinson 2002; Carrapiço 2006).

SA can be synthesized via two distinct routes in land plants: In *Arabidopsis thaliana* the majority of SA stems from conversion of chorismate into isochorismate by isochorismate synthase (ICS) (Wildermuth et al. 2001, Garcion et al. 2008). Isochorismate is transferred from the chloroplast to the cytoplasm and as a substrate of avrPphB SUSCEPTIBLE3

(PBS3) conjugated with glutamate, resulting in the product isochorismate-9-glutamate (Rekhter et al. 2019). A spontaneous reaction, which can be also catalysed by ENHANCED PSEUDOMONAS SUSCEPTIBILTY 1 (EPS1), results in the conversion of isochorismate-9-glutamate to SA (Rekhter et al. 2019, Torrens-Spence et al. 2019). Yet, *EPS1* appears to be specific to Brassicaceae and a clear *PBS3* ortholog has not been identified in *A. filiculoides* (de Vries et al. 2018, Torrens-Spence et al. 2019, Li et al. 2020).

Already in other angiosperms than *A. thaliana*, SA may not come from the ICS-dependent pathway but rather derives from benzoic acid (Meulwy et al. 1995, Pallas et al. 1996, Coquoz et al. 1998). These early publications suggested that SA is synthesized via a Phenylaline Ammonia Lyase (PAL)-dependent pathway from phenylalanine. Phenylalanine is converted into *trans*-cinnamic acid (Widhalm and Dudareva 2015). From here, SA can be synthesized via peroxisomal β-oxidation or two cytoplasm-localized non-oxidative pathways, one CoA-dependent and one CoA-independent (Widhalm and Dudareva 2015). None of the pathways has so far full enzymatic evidence. Yet, some steps along the way have been functionally characterized. Some first importers of phenylpropanoid pathway-derived compounds into the peroxisome have been identified (Block et al., 2014). In the oxidative peroxisomal pathway, *trans*-cinnamic acid is converted into cinnamoyl-CoA (if the latter is not imported) via *trans*-cinnamic acid ligase (*Ph-*CNL) from *Petunia hybrida* and possibly also by its ortholog BENZOYLOXYGLUCOSINOLATE 1 (BZO1) in *A. thaliana* (Colquhoun et al. 2012, Klempien et al. 2012, Lee et al. 2012). In *P. hybrida* the following steps towards benzoyl-CoA are catalyzed via chalcone dehydrogenase (*Ph*CHD) and katalase1 (*Ph*KAT1) (Van Moerkercke et al. 2009, Qualley et al. 2012). In *A. thaliana* they are suggested to be realized by the multifunctional β-oxidation enzyme abnormal inflorescence meristem1 (*At*AIM1) and *At*KAT2 (Bussell et al. 2014, Widhalm and Dudareva 2015). The resulting benzoyl-CoA is converted to benzoic acid by a thioesterase, likely 1,4-dihydroxy-2-naphthoyl (DHNA)-CoA THIOESTERASE 1 (*At*DHNAT1/2) in *A. thaliana* (Widhalm et al. 2012, Widhalm and Dudareva 2015) or Ph-TE1 in *P. hybrida* (Adebesin et al. 2018). In the non-oxidative cytoplasmic pathways, a hydratase is suggested to use *trans-*cinnamic acid or cinnamoyl-CoA as a substrate and hydrates the double bond to form 3-hydroxy-3-phenylpropionic acid or 3-hydroxy-3-phenylpropanoyl-CoA, respectively (see Widhalm and Dudareva, 2015). Both of which are likely converted into benzaldehyde by a lyase-dependent reaction (see Widhalm and Dudareva, 2015). In *A. thaliana* benzaldehyde is converted to benzoic acid by Arabidopsis Aldehyde Oxidase4 (AAO4) (Ibdah et al. 2009), while in Snapdragon (*Antirrhinum majus*) it is imported into the mitochondrion and converted to benzoic acid by benzaldehyde dehydrogenase (BALDH) (Long et al. 2009). Independent of the route benzoic acid is produced, it was suggested that benzoic acid 2-hydroxylase (BA2H)-like enzyme catalyzes the last step from there to SA (León et al. 1995).

In this study we explore the connections between SA and the coordinated symbiosis of the water fern *A. filiculoides* and its cyanobiont *T. azollae*. Using HPLC-MS/MS we measured SA in roots and the photosynthesizing sporophyte body of *A. filiculoides*. Because it is not known by which pathway the symbiotic system synthesizes SA, we used the cumulated knowledge on enzymatic synthesis of SA of the green lineage to identify candidate genes for SA biosynthesis in *A. filiculoides*. Applying comparative genomics and phylogenetics, we searched for orthologs—and the closest homologs—of key genes likely acting in the ICS- and PAL-dependent pathway; we further explored the possibility of cyanobiont-derived SA. We find that only the genomes of *Chara braunii, Selaginella moellendorffii* and angiosperms encode a separate standalone ICS. All other chlorophyte and streptophyte algae as well as bryophytes and the fern *Salvinia cucullata* contain a fusion enzyme, where the ICS domain is fused with a MenC and/or MenD domain, suggesting that chorismate can be quickly diverted into the phylloquinone pathway. Moreover, *A. filiculoides* entirely lacks an ICS-encoding gene. In contrast, our phylogenetic data support the presence of homologs for all known steps in the PAL-dependent pathway across land plants, suggesting that the benzoic acid pathway has the potential to be recruited for SA biosynthesis in several land plant lineages. Our data pinpoint the biosynthetic routes of SA via benzoic acid as the most likely origin of SA in *A. filiculoides*. We carried out global gene expression analyses on the *PAL*-dependent SA biosynthesis pathway and homologs to SA responsive genes. The majority of SA-associated genes that showed differential gene expression patterns appear to be down-regulated in the presence of the cyanobionts. Integrating this with previous results, we hypothesize that a feedback loop has co-evolved between the two partners, that tightly co-ordinates the levels of SA with the proliferation of the cyanobionts.

## MATERIAL AND METHODS

### Measurement of SA levels

The water fern *Azolla filiculoides* was cultured at 75% relative humidity in a heat-sterilized glass vessel. The ferns floated on 250 mL filtered water with a pH of 7.0. During the 16h-long day, the fern was exposed to 450 µmol quanta m^-2^ s^-1^ at 24 °C; during the 8h-long night, the temperature dropped to 20 °C. We picked plants from actively growing culture and used sterilized micro-scissors to separate the fern body into whole roots and green sporophyte tissue (i.e. with roots removed). Extraction was performed as previously described for lipids (Matyash et al., 2008), with some modifications specified previously (Iven et al., 2012). The analysis was performed using an Agilent 1100 HPLC system (Agilent, Waldbronn, Germany) coupled to an Applied Biosystems 3200 hybrid triple quadrupole/linear ion trap mass spectrometer (MDS Sciex, Ontario, Canada) by using an ESI chip ion source (TriVersa NanoMate; Advion BioSciences, Ithaca, NY, USA). The quantification of SA was performed as previously described (Iven et al., 2012) applying a scheduled multiple reaction monitoring detection in negative ionisation mode including the following MS transitions: 141/97 [declustering potential (DP) −45 V, entrance potential (EP) −7 V, collision energy (CE) −22 V] for D_4_-SA and 137/93 (DP −45 V, EP −7 V, CE −22 V) for SA. The quantification was carried out using a calibration curve of intensity (m/z) ratios of [SA]/[D_4_-SA] vs. molar amounts of SA (0.3-1000 pmol).

Statistical analyses have been performed in R v.3.6.0. Data was tested for normality using a Shapiro-Wilks test (Shapiro and Wilk, 1965), following a test to compare the variance of the data between replicates of SA measurements from roots and sporophytes of *A. filiculoides*. The data was normally distributed and showed no significant difference in variance. Accordingly, a two-sample t-test (Student 1908) was computed.

### Identification and phylogenetic analyses of SA biosynthesis genes of *A. filiculoides*

For phylogenetic analyses, we worked with protein data from the genomes of *Anthoceros agrestis* as well as *Anthoceros punctatus* (Li et al., 2020), *Amborella trichopoda* (*Amborella* Genome Project, 2013), *Arabidopsis thaliana* (Lamesch et al., 2011), *Azolla filiculoides* (Li et al., 2018), *Brachypodium distachyon* (The International Brachypodium Initiative, 2010), *Capsella grandiflora* (Slotte et al., 2013), *Gnetum montanum* (Wan et al., 2018), *Marchantia polymorpha* (Bowman et al., 2017), *Nicotiana tabacum* (Sierro et al., 2014), *Oryza sativa* (Ouyang et al., 2007), *Picea abies* (Nystedt et al., 2013), *Physcomitrium patens* (Lang et al., 2018), *Salvinia cucullata* (Li et al., 2018), *Selaginella moellendorffii* (Banks et al., 2011), and *Theobroma cacao* (Argout et al., 2011); (b) the genomes of seven streptophyte algae: *Chlorokybus atmophyticus* (Wang et al., 2020), *Chara braunii* (Nishiyama et al., 2018), *Klebsormidium nitens* (Hori et al., 2014), *Mesotaenium endlicherianum* (Cheng et al., 2019), *Mesostigma viride* (Wang et al., 2020), *Penium margaritaceum* (Jiao et al., 2020), *Spirogloea muscicola* (Cheng et al., 2019). For each of the protein families, specific cutoffs were chosen (see below). All sequences that have met the respective cutoffs, were aligned using MAFFT v7.453 (Katoh and Standley, 2013) with the setting L-INS-I. Alignments were cropped to conserved and alignable regions for all homologs. Maximum likelihood phylogenies were computed using IQ-TREE multicore v.1.5.5 (Nguyen et al., 2015) with 100 bootstrap replicates. For finding the best model of protein evolution, we used ModelFinder (Kalyaanamoorthy et al., 2017). The best models were LG+G4 (Le and Gascuel, 2008) for PAL, LG+I+G4 for AIM1, KAT, LG+F+I+G4 for AAO, JTT+I+G4 (Jones et al., 1992) for DHNAT.

Protein sequences for enzymes involved in PAL-dependent biosynthesis of SA in *A. thaliana, P. hybrida* or Snapdragon (*Antirrhinum majus*) have been downloaded from TAIR (Lamesch et al., 2011), NCBI, and UniProt. Sequences have sampled by using an HMM search via HMMER 3.1b2 (Mistry et al., 2013) with the ammonia lyase domain (obtained from PFAM) as a query against the proteins from the above-described genomes; all sequences that met the inclusion threshold were retained. Additionally, HAL and fungal PAL sequences that were included in de Vries et al. (2017) were added. For computing the maximum likelihood phylogeny, we used only those sequences that were at least 600 amino acids in length.

For identification of putative ICS sequences in *A. filiculoides*, we conducted an HMM search using HMMER 3.1b2 (Mistry et al., 2013) for the chorismate binding domain against the high- and low-confidence protein sequences from the *A. filiculoides* genome v.1.1 (Li et al., 2018). We used all three thusly identified *Azolla* sequences that met the HMM search inclusion threshold (*e* values of 4.8*10^−87^, 4.6*10^−85^, and 3.7*10^−84^) as queries for a BLASTp search against the protein dataset from the above described genomes. To improve the sampling on the closest algal relatives of land plants, we additionally included sequences found via a tBLASTn in the transcriptomes of *Spirogyra pratensis* (de Vries et al., 2020), *Zygnema circumcarinatum* (de Vries et al., 2018), and *Coleochaete orbicularis* (Ju et al., 2015). We used only those sequences that have met an e value cutoff of 10^−10^, had a minimum length of 300 amino acids.

### Identification of SA biosynthesis genes in cyanobacteria

To identify the ICS of *T. azollae* we downloaded the ICS sequence of *Trichormus variabilis* from NCBI and used it in a BLASTp query against *T. azollae* in the non-redundant protein (nr) database of NCBI. To identify chorismate binding enzymes we blasted ICS, anthranilate synthase and aminodeoxychorismate synthase of *T. azollae* against cyanobacteria in the nr database. Further, we downloaded sequences for isochorismate pyruvate-lyase (IPL) from *Pseudomonas aeruginosa* (accession NP_252920.1) and salicylate synthase (SAS) of *Mycobacterium tuberculosis* (accession YP_177877.1). We used these sequences to identify IPL and SAS enzymes in cyanobacteria using BLASTp at NCBI.

### Identification of homologs to SA responsive genes

We downloaded all protein sequences from TAIR10 (Lamesch et al. 2011) linked to the term response to SA. This resulted in 176 sequences, which we used as a query in a blastp search against the proteome of *A. filiculoides* (Li et al. 2018). The 16,776 non-unique hits were funneled into a reciprocal BLASTp search with max_target_seqs 1, returning the best hits from *A. thaliana* for 3,386 unique *A. filiculoides* proteins. We next retained only those sequences that had their best hit to one of the 176 accessions of *A. thaliana* associated with the term response to SA and used an e-value cutoff of 10^−7^. The *A. filiculoides* sequences corresponding to the final list of hits against *A. thaliana* were considered homologs to SA responsive genes; and after removal of low-confidence proteins added up to 97 candidates.

### Analyses of conserved sites in chorismate binding enzymes

Protein sequences of the chorismate binding enzymes of plants were separated into ICS, anthranilate synthase and aminodeoxychorismate synthase sequences according to the phylogenetic analysis. Further protein sequences of the top 100 blastp hits for each type of the cyanobacterial chorismate binding enzymes were downloaded from NCBI. Each subset of protein sequences was aligned with that of the anthranilate synthase of *Salmonella enterica* subsp. *enterica* serovar Typhimurium str. LT2 with MAFFT using a L-INS-I approach (Katoh and Standley, 2013) to identify the conserved positions described in Plach et al. (2015). To generate sequence logos for the respective groups the sequence from *Salmonella* was removed from the alignment after the positions were located. Sequence logos were generated based on the alignments.

### Transcriptomic profiling of candidate genes for SA biosynthesis and signaling

Raw read count data for *A. filiculoides* with or without cyanobiont and treated with or without fixed nitrogen (NH_4_NO_3_) supply were obtained from Eily et al. (2019). We calculated the TPM for all datasets according to (Robinson and Oshlack, 2010) and used edgeR (Robinson et al., 2010) to calculate log_2_-foldchange and to identify the differently expressed genes (DEGs) within a given dataset. To calculate FDR from p-values a Benjamini-Hochberg correction was used.

### Protein domain search

We used Interpro (Blum et al., 2020) to identify protein domains in chorismate binding enzymes of land plants and used the information of TIGR to identify the putative function of the chorismate binding enzymes identified in our BLASTp and phylogenetic analyses. Likewise, we used CD search (Marchler-Bauer et al., 2004) and the information of TIGR IDs for protein domain analyses of cyanobacterial protein domains of chorismate binding enzymes.

## RESULTS AND DISCUSSION

### *Azolla filiculoides* produces SA

Molecular data on cross-talk and co-ordination between *Azolla* and its cyanobionts are just becoming more and more abundant (Brouwer et al. 2017, de Vries et al. 2018, Li et al. 2018, Eily et al. 2019, Güngör et al. 2020). Several studies point to a role of flavonoids in the communication between the two partners (Pereira and Carrapiço 2007, Güngör et al. 2020) and MeSA, a methylated, volatile derivative of SA, appears to also interfere in the interaction (de Vries et al. 2018). Yet, we do not know whether and how SA is synthesized endogenously in *A. filiculoides*.

In a first step, we measured endogenous SA levels in the roots and sporophyte of *A. filiculoides* (Table 1). The data confirmed that *A. filiculoides* harbors SA. Roots appear to harbor slightly more SA than the sporophyte (roots: 0.18±0.03 nmol/g fresh weight (FW), sporophyte: 0.09±0.02 nmol/g FW, p-value=0.0337). Yet, these basal levels of SA in the non-axenic *Azolla* are relatively low compared to for example Col-0 from *A. thaliana* (Rekhter et al. 2019). Regardless of how much SA was measured, the mere fact that it is present in the fern body begs the question of how it is synthesized. We thus next used the recently released genomic data on *Azolla filiculoides* to explore possible biosynthetic routes towards SA.

**Table 1.** Quantification of salicylic acid (SA) in root and green sporophyte tissues of *Azolla filiculoides* via HPLC-MS/MS.

| tissue type | biological replicate | weight f.w. [g] | SA [nmol/g f.w.] |
| --- | --- | --- | --- |
| whole root | I | 0.02 | 0.22 |
|  | II | 0.01 | 0.17 |
|  | III | 0.025 | 0.14 |
| sporophyte | I | 0.105 | 0.1 |
|  | II | 0.1 | 0.11 |
|  | III | 0.11 | 0.07 |

Abbreviations: f.w. = fresh weight

### A treatise of the three possible routes for SA biosynthesis in *A. filiculoides*

Land plants have two biosynthetic routes towards SA. One route depends on ICS, the other on PAL (Vlot et al. 2009). Unlike other land plants, *Azolla* theoretically has a third option to acquire SA via the biochemical capacities of its omnipresent cyanobiont. Here we used a phylogenetic approach utilizing the genome data of *A. filiculoides* and a diversity of genomes from other species from the green lineage to identify putative candidates for the SA biosynthesis pathways in *Azolla*. We further investigated the cyanobiont genome with regard to its theoretical ability to synthesize the hormone.

#### The ICS-dependent pathway

The ICS-pathway is localized in the plastid and cytoplasm and requires ICS1 and 2 and PBS3 (Wildermuth et al. 2001, Garcion et al. 2008, Rekhter et al. 2019). The last step in the conversion to SA can either be spontaneous or catalyzed by EPS1 (Rekhter et al. 2019, Torrens-Spence et al. 2019). EPS1 is Brassicaceae-specific (Torrens-Spence et al. 2019), but it is also not strictly required for the interaction. PBS3 has emerged from a lineage-specific expansion of GH3-encoding genes (de Vries et al. 2018, Li et al. 2020). However, two GH3-homologs exist in the *A. filiculoides* genome that fall into the same GH3 clade as *At*PBS3 (Li et al. 2018, Li et al. 2020 supplemental material). For the key enzyme, ICS, BLASTp surveys identified only a single low-confidence sequence in *A. filiculoides* (Li et al. 2018, 2020), which appears rather divergent to other ICS1 candidates as shown in the (supplemental) phylogenetic analyses included in Li et al. (2020). Here, we investigated the chorismate binding protein families in more detail.

Using an HMMsearch against the genome of *A. filiculioides*, we found three protein-coding genes (*Azfi_s0185*.*g056617, Azfi_s0002*.*g007267*, and *Azfi_s0061*.*g034905*) with a chorismate-binding domain (for which we obtained an HMM profile from PFAM; El-Gebali et al., 2019). ICS sequences exist throughout the green lineage, but we found no ortholog for *A. filiculoides* (Figure 1a). The existence of an ICS sequence in land plants and algae suggests that the last common ancestor of the Chloroplastida had an ICS sequence. The lack of an ICS in *A. filiculoides* can only be explained by a secondary loss; this absence is corroborated by the lack of clear ICS orthologs in the transcriptome data of *Azolla carolinia* (sequenced in the framework of the 1KP efforts; Carpenter et al., 2019) and *Azolla pinnata* (Shen et al., 2018; see supplemental Figure S1).

**Figure 1.**
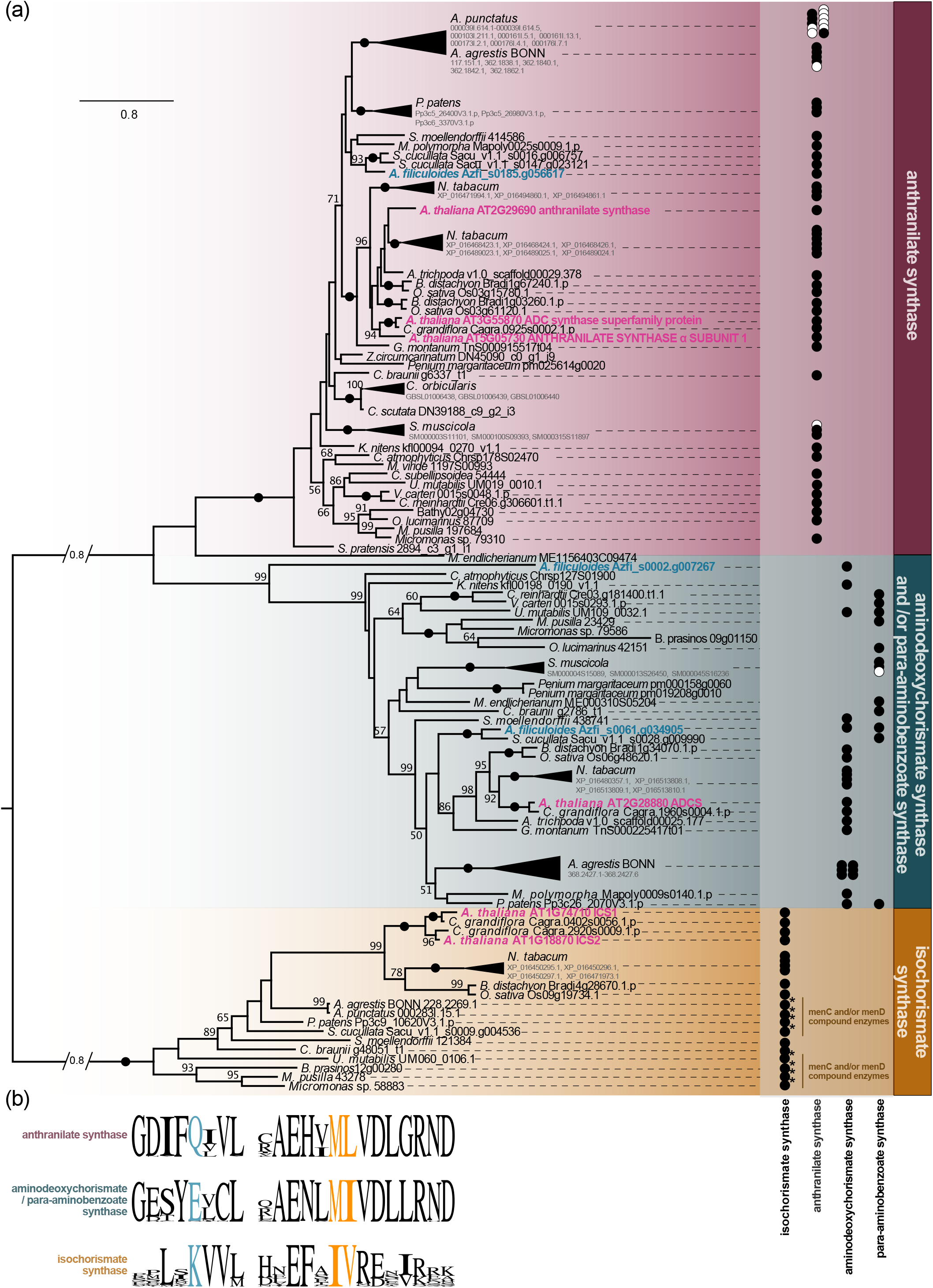
Evolution of chorismate binding enzymes in the green lineage. (a) ML-phylogeny of chorismate binding enzymes from Chloroplastida with 100 bootstrap replicates. Sequences from *A. thaliana* are highlighted in pink and sequences of *A. filiculoides* are highlighted in blue. We recovered three clear clades, one for isochorismate synthases (ICS), one for aminodeoxychorismate synthases and *p*-aminobenzoate synthases and one for anthranilate synthases. Bootstrap support of 100 is indicated by black dots on the branches. On the right we indicate TIGR domain annotations for the for different categories of chorismate binding enzymes. Black dots indicate which domain was found; no dot indicates that no TIGR domain annotation was present for this sequence. White dots only occur for collapsed branches, to indicate for which of the collapsed sequences a domain annotation was available (black dot) and for which not (white dot). (b) Conserved sequence logos for the clades containing anthranilate synthase, aminodeoxychorismate/*p*-aminobenzoate synthase and isochorismate synthase (ICS) shown in (a). Highlighted residues correspond to amino acids that have been suggested to be functionally relevant for the product specificity of these enzyme categories in Plach et al. (2015).

Whether the isochorismate pathway is used in all species with an ICS ortholog is however unclear when we investigated the protein domain structures of all ICS sequences. The ICS candidates from bryophytes, algae and the fern *S. cucullata* appear to be compound enzymes coupled with a MenC and/or MenD domain (Figure 1a). This suggests that in these species ICS may directly funnel isochorismate into the menaquinone pathway rather than into a route towards SA. Yet, in some of the species, such as the moss *Physcomitrium patens*, SA has been detected (Ponce De León et al., 2012). Thus, ICS was either recruited later in the evolution of land plants as the primary route towards SA or it is species specific whether SA derives from isochorismate or benzoic acid in plants. However, ICS may not be the only source for isochorismate.

Usually anthranilate synthases convert chorismate to aminodeoxyisochorismate and then to anthranilate. That said, a *Salmonella* anthranilate synthase was successfully engineered to become an isochorismate-forming enzyme by exchanging two amino acids in the active center (Plach et al. 2015). A mutation leading to a lysine at position 263 instead of glutamine and another either at 364 changing a methionine to a lysine or at 365 changing lysine to valine led to isochorismate formation in *Salmonella* (Plach et al. 2015). These residues are more or less conserved across different phyla in bacteria (Plach et al. 2015): with Q263 and M364 being conserved across all analyzed anthranilate and aminodeoxychorismate synthases and L365 in the anthranilate synthases and I365 in the aminodeoxychorismate synthases. ICS and salicylate synthase (SAS) show a conserved K263, but appeared to be slightly more variable at positions 364 (ICS major amino acid (aa): 364L; SAS major aa: 364I) and position 365 (ICS major aa: 365V; SAS major aa: 365S). Further, the 263 lysine (K190 in *Escherichia coli*) was shown to be highly relevant for the function of bacterial ICS (Kolappan et al. 2007). We found that all land plant lineages included in this phylogeny, including *A. filiculoides*, also encode putative anthranilate synthases in their genomes (Figure 1a).

To elucidate whether the putative anthranilate synthases of Chloroplastida and of *A. filiucloides* in particular can potentially take over the role of ICS in the fern, we explored the respective positions determining the product specificity (a) in the set of land plant chorismate binding enzymes to establish the most abundant residues in land plants at these functionally relevant positions and (b) in the three *A. filiculoides* protein sequences. The anthranilate synthases of plants had the same aa pattern as bacterial anthranilate synthases: Q, M, and L (Figure 1b), while the aminodeoxychorismate synthases encoded mainly an E at the position equivalent to 263, and M, and I at the respective positions for 364 and 365 (Figure 1b). 263E was the second most abundant residue in aminodeoxychorismate synthases of bacteria (Plach et al. 2015), suggesting that the three residues are in general conserved across prokaryotes and eukaryotes. Plant ICS however showed a K, I and V at the respective positions in the alignment. All three positions were not variable in plant ICS (Figure 1b). This is corroborated by the fact, that the anthranilate synthase from *A. filiculoides* encoded a QML motif as any other land plant. Overall, this suggests that it does not fill in for the function of the missing ICS.

#### The PAL-dependent pathway

A PAL-dependent pathway for SA biosynthesis has been suggested for several angiosperms (Meuwly et al. 1995, Pallas et al. 1996, Coquoz et al. 1998). The synthesis of SA via PAL can be facilitated via several different routes of benzoic acid metabolism. All of them have in common that PAL, which converts phenylalanine to *trans*-cinnamic acid, is the entry point for SA synthesis (Widhalm and Dudareva, 2015). The β-oxidation pathway requires enzymes from the 4CL family and the closely related *At*BZO1/*Ph*-CNL clade, the hydratase *Ph*CHD, *Ph*KAT1and *At*DHNAT1/2 (Van Moerkercke et al., 2009, Colquhoun et al., 2012, Klempien et al., 2012, Lee et al., 2012, Qualley et al., 2012, Widhalm et al., 2012, Widhalm and Dudareva, 2015). The non-oxidative pathway also requires 4CL, the hydratase AIM1, which is the ortholog of *A. thaliana* to *Ph*CHD, a lyase, AAO4 (Ibdah et al., 2009, Bussel et al., 2014, Widhalm and Dudareva, 2015) or in Snapdragon (*Antirrhinum majus*) the benzaldehyde dehydrogenase (BALDH; Long et al. 2009). The last step in the biosynthesis of SA is only indirectly characterized and an inhibitor study suggests that BA2H-like enzyme is required (León et al., 1995).

To identify PAL candidates in *A. filiculoides*, we performed an HMMsearch with the aromatic lyase motif (HMM profile obtained from PFAM; El-Gebali et al., 2019) against the genome of *A. filiculoides* as well as other representative species from across the streptophyte tree of life. The aromatic lyase motif is present in PAL, (P)TAL, TAM and PAL-tRNA synthase fusion proteins of plants. We used this dataset together with a subset of fungal PAL and eukaryotic HAL sequences (selected from the dataset of de Vries et al. 2017) to build a phylogeny.

In our phylogeny the fungal PAL, tRNA-PAL fusion proteins and eukaryotic HAL proteins each form their own clade (Figure 2). Yet, the functionally characterized PTAL and TAM sequences of plants fall into the midst of the putative PAL clade (Figure 2). A detailed study in *Sorghum bicolor* has identified two positions relevant for distinguishing PAL and PTAL (Jun et al. 2018): the presence of Phe at position 123 (relative to the characterized *Sb*PTAL Sb04g026510) and Lys at position 443, were present in sequences with PAL activity, while *Sb*PTALs showed His and Lys at the two respective positions. In *Os*TAM (Yan et al. 2015) these two positions are occupied by a Tyr and Asn respectively; the same combination of residues occurs in a *S. bicolor* PAL/PTAL candidate (Figure 2) that had neither tyrosine nor phenylalanine activity (Jun et al. 2018). To make more sense of the phylogenetic data, we mapped this sequence information of the two positions to the phylogeny.

**Figure 2.**
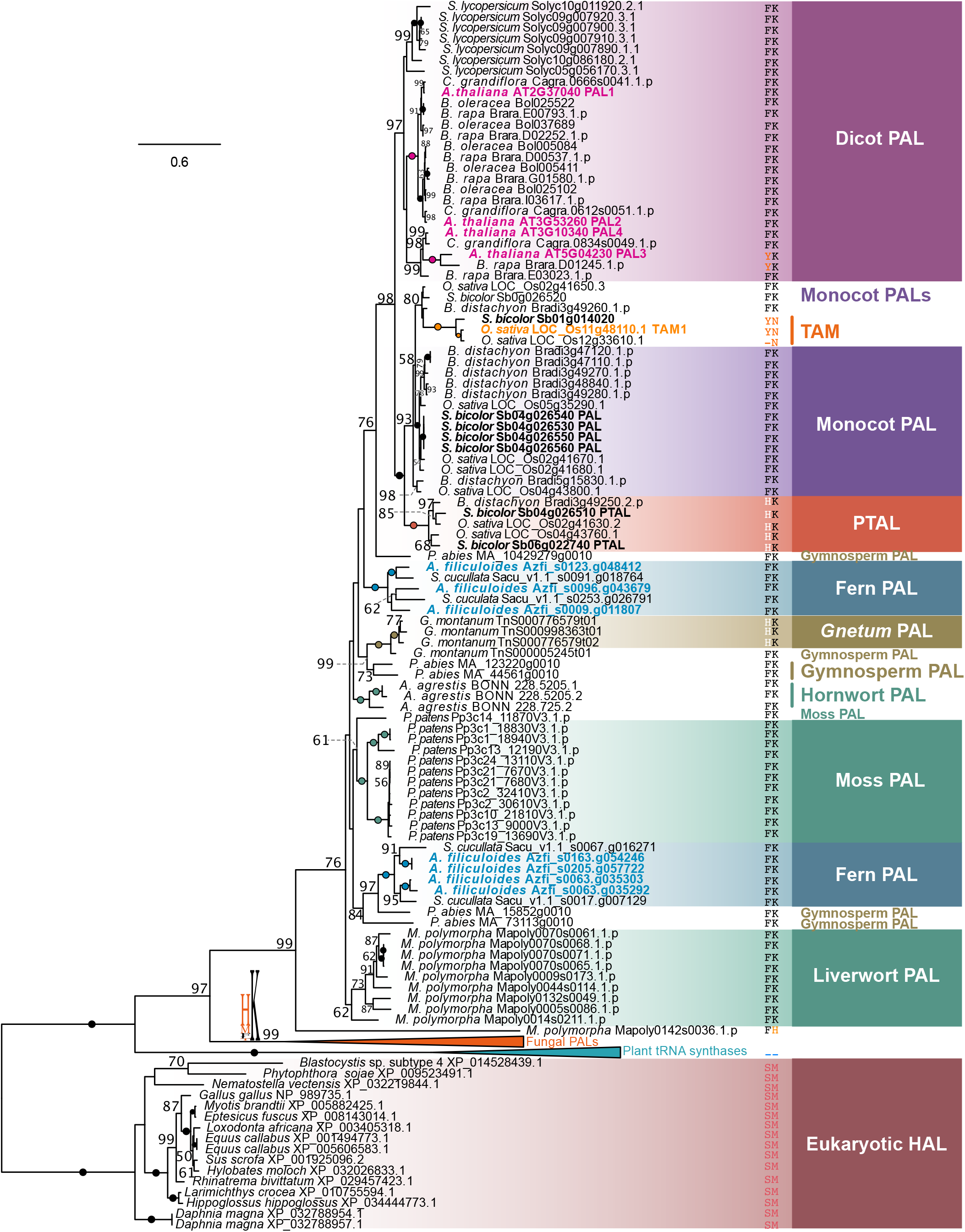
Phylogeny of PAL and HAL sequences from eukaryotes and prokaryotes. ML-phylogeny of eukaryotic phenylammonia lyases (PAL) and HALs with 100 bootstrap replicates; LG+G4 was chosen as model for protein evolution according to Bayesian Information Criterion. Sequences for PAL in plants have been identified using an HMM approach. Sequences from fungal PALs and eukaryote HALs have been selected from de Vries et al. (2017). Sequences from *A. thaliana* have been highlighted in pink, *A. filiculoides* is highlighted in blue, sequences from *S. bicolor* are indicated in black and TAM1 from *Oryza sativa* is highlighted in orange. Bootstrap support of 100 is indicated by black and colored dots on the branches. The clades with fungal PAL and plant tRNA synthase sequences have been collapsed. The substrate-specificity defining residues 123 and 443 of *S. bicolor* PTAL (Sb04g026510) have been analyzed for all sequences in the alignment and are indicated on the right next to the respective sequences. One exception is the collapsed branch of fungal PALs. Due to the variance of residue 123 the sequence logo for the two residues is printed on the branch itself.

While all HAL sequences occupied a 123S and 443M, the tRNA synthesis fusion proteins showed a deletion of both residues. In contrast, most sequences that fall into the plant and fungal PAL clade occupy Lys at position 443 (relative to Sb04g026510), with the exception of one very divergent sequence from *M. polymorpha* (Mapoly0142s0036.1.p; 443H) and the subclade including *Os*TAM (LOC_Os11g48110.1), which showed Asn at position 443 (Figure 2). Position 123 was more variant among fungal and plant PALs: Most fungal PALs have His at the relevant site, suggesting they may be PTALs instead. In contrast, most plant PAL candidates have a Phe at the site of interest, with the exception of the subclades including the monocot PTALs and TAM candidates and one *A. thaliana* PAL sequence (123Y in *At*PAL3; Figure 2). In agreement, *At*PAL3 was found to have the lowest catalytic activity as phenylalanine ammonia lyase of all four PALs of *A. thaliana* (Cochrane et al. 2004). All seven included sequences of *A. filiculoides*, that were full-length and high-confidence sequences, showed a 123F and 443K (Figure 2), suggesting them as likely PAL candidates.

Next, we investigated the acyl-activating enzyme (AAE) family that includes *Ph*-CNL and its co-ortholog *At*BZO1 (Figure S2), and all *At*4CLs. Our analysis captured sequences from five of the seven predicted subfamilies of AAEs (Shockey et al., 2003). In general, we recovered four of the AAE subfamilies with a bootstrap support of at least 75. The family shows various lineage-specific expansions (Figure S2). Only AAE clade VII did not form a monophyletic clade (Figure S2). Clade I, IV, V and VI appear to have their origin in the last common ancestor of land plants, yet especially clade IV to VI show substantial lineage-specific radiation. Our enzymes of interest belong to these highly expanded clades: *At*BZO1 and *Ph*-CNL belong to AAE clade VI, and 4CL1,2,3 and 5 belong to AAE clade IV (Figure S2, Shockey et al., 2003). Despite that, we found four clear homologs of *A. filiculoides* for the 4CL-encoding genes (Azfi_s0003.g007625, Azfi_s0013.g013344, Azfi_s0114.g046013 and Azfi_s0030.g024259) and two candidates for AAE clade VI (BZO1/*Ph*-CNL; Azfi_s0159.g053935 and Azfi_s0002.g003721 [low-confidence sequence]). The next steps in the oxidative and non-oxidative pathways are catalyzed by hydratases *Ph*CHD in *P. hybrida* and likely by its ortholog *At*AIM1 in *A. thaliana* (Qualley et al. 2012, Bussell et al. 2014). Our phylogeny shows that an AIM1/CHD hydratase was present in the last common ancestor of Chloroplastida. While several lineages have species-specific expansions of the AIM1/CHD family, *A. filiculoides* possesses one clear ortholog for this hydratase (Azfi_s0256.g060521; Figure 3). A similar picture emerges for the next step in the peroxisomal pathway catalyzed by KAT: One KAT homolog was present in the last common ancestor of Chloroplastida, but KAT sequences have expanded in several lineages, most pronounced in the Brassicaceae (Figure 4). In contrast, the two sequenced fern genomes (*S. cucullata* and *A. filiculoides*), however, possess only one copy for KAT (Azfi_s0001.g000824, Figure 4). The final peroxisomal step to benzoic acid is realized by a thioesterase; *At*DHNAT1/2 shows activity on benzoyl-CoA esters (Widhalm et al. 2012) and was hypothesized for this step (Widhalm and Dudareva 2015). The DHNAT family more or less mirrors the species phylogeny, yet includes again species-specific radiations (Figure 5). In the ferns, we find a duplication for DHNAT homologs in *S. cucullata*, but one ortholog in *A. filiculoides* (Azfi_s0008.g011631; Figure 5).

**Figure 3.**
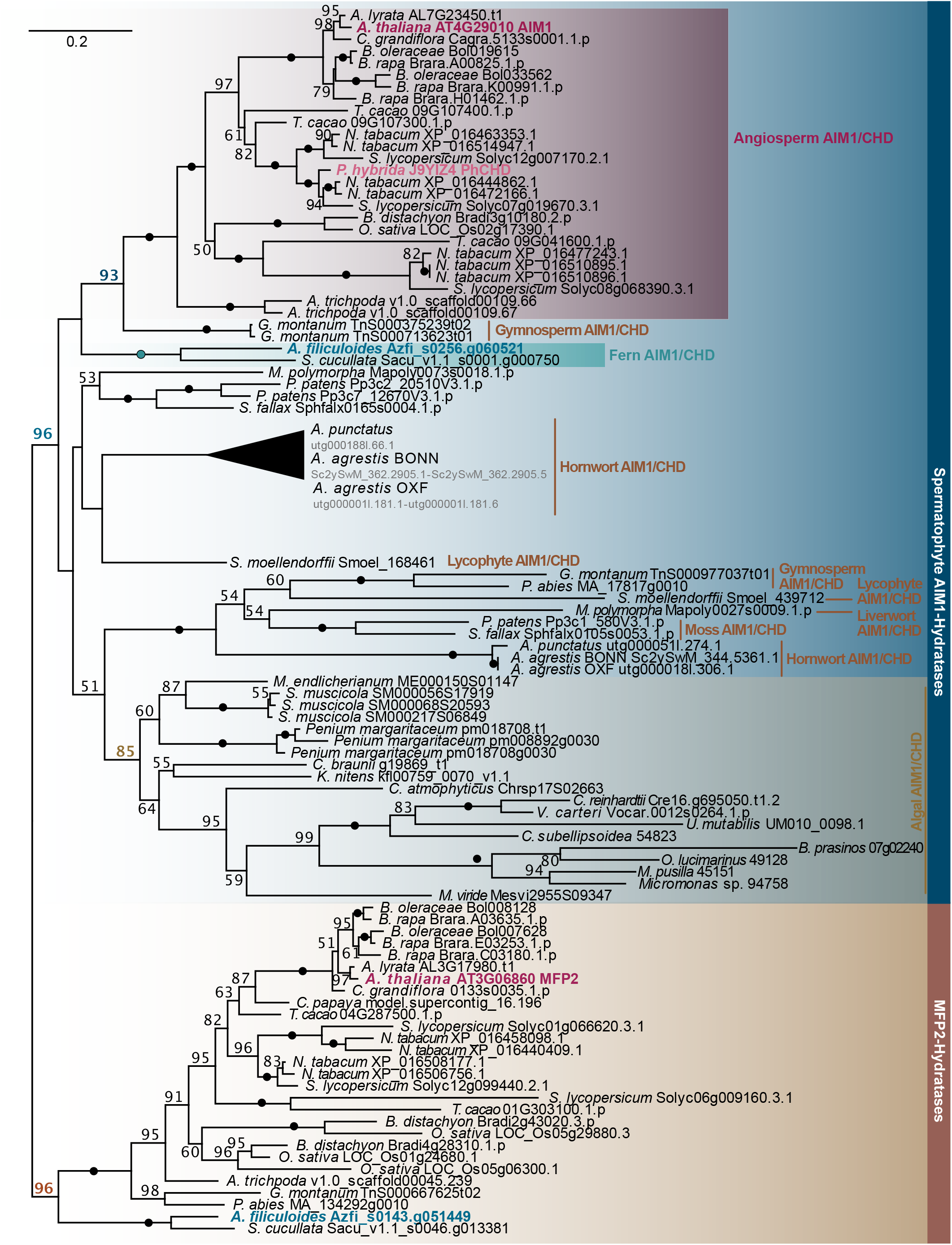
Phylogeny of Hydratases in Chloroplastida. ML-phylogeny pointing the evolutionary history of AIM1/CHD hydratases in chloroplastida using 100 bootstrap replicates; LG+I+G4 was chosen as model for protein evolution according to Bayesian Information Criterion. The AIM1 sequence of *A. thaliana* is highlighted in purple, its ortholog *Ph*CHD from *Petunia hybrida* is highlighted in light pink and sequence from *A. filiculoides* is highlighted in blue. Bootstrap levels = 100 are indicated by black dots.

**Figure 4.**
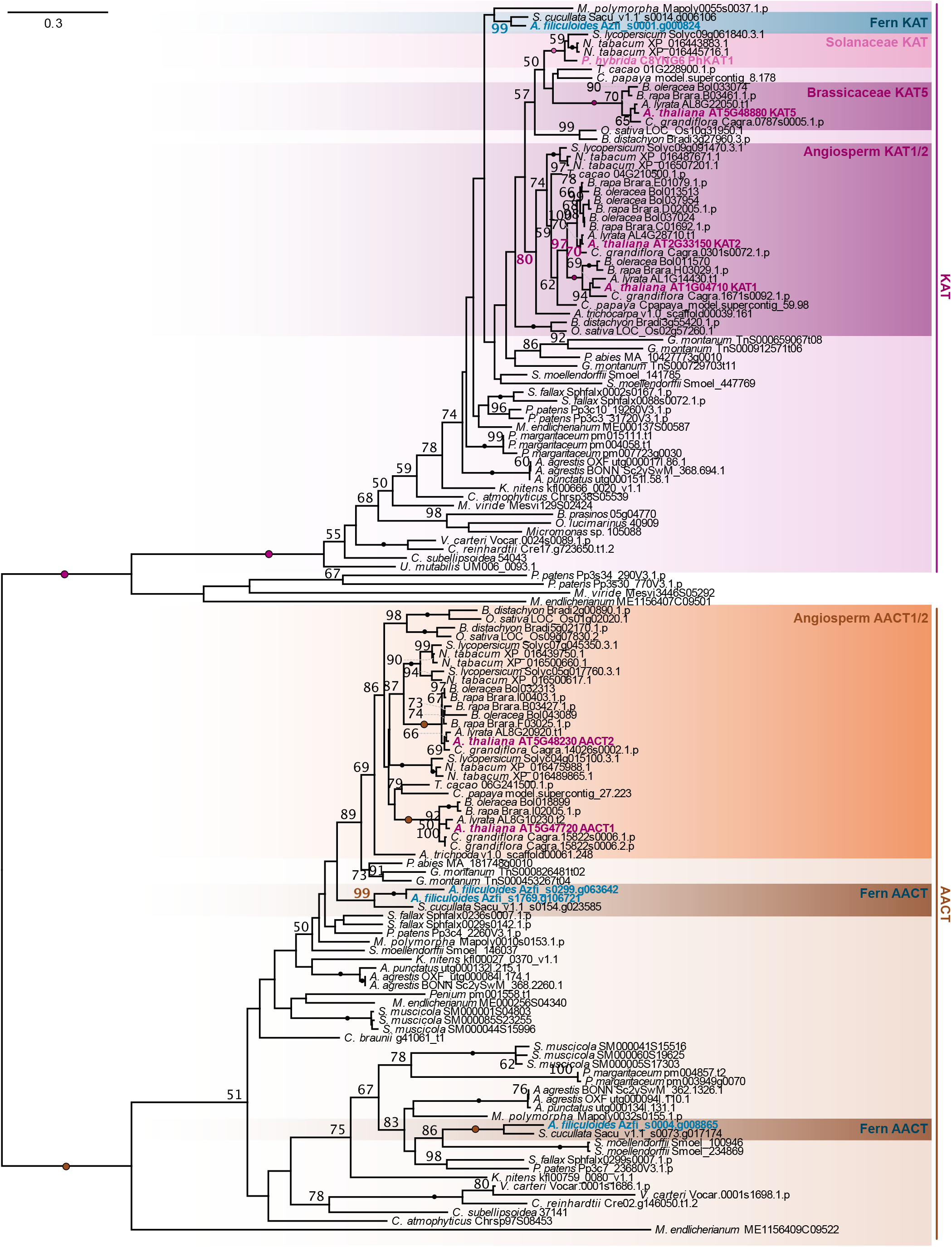
ML-phylogeny determining the evolutionary history of KAT sequences in chloroplastida. The phylogeny is based on 100 bootstrap replicates; LG+I+G4 was chosen as model for protein evolution according to Bayesian Information Criterion. Sequences from *A. thaliana* are shown in purple, the *Ph*KAT sequence from *P. hybrida* is shown in light pink and the sequence from *A. filiculoides* is highlighted in blue. Clades with brassicaceae and angiosperm KAT sequences have been highlighted in purple, Solanaceae KAT sequences are indicated by a pink box and fern KAT sequences are indicated by a blue box. Bootstrap levels of a 100 are indicated by dots along the branches.

**Figure 5.**
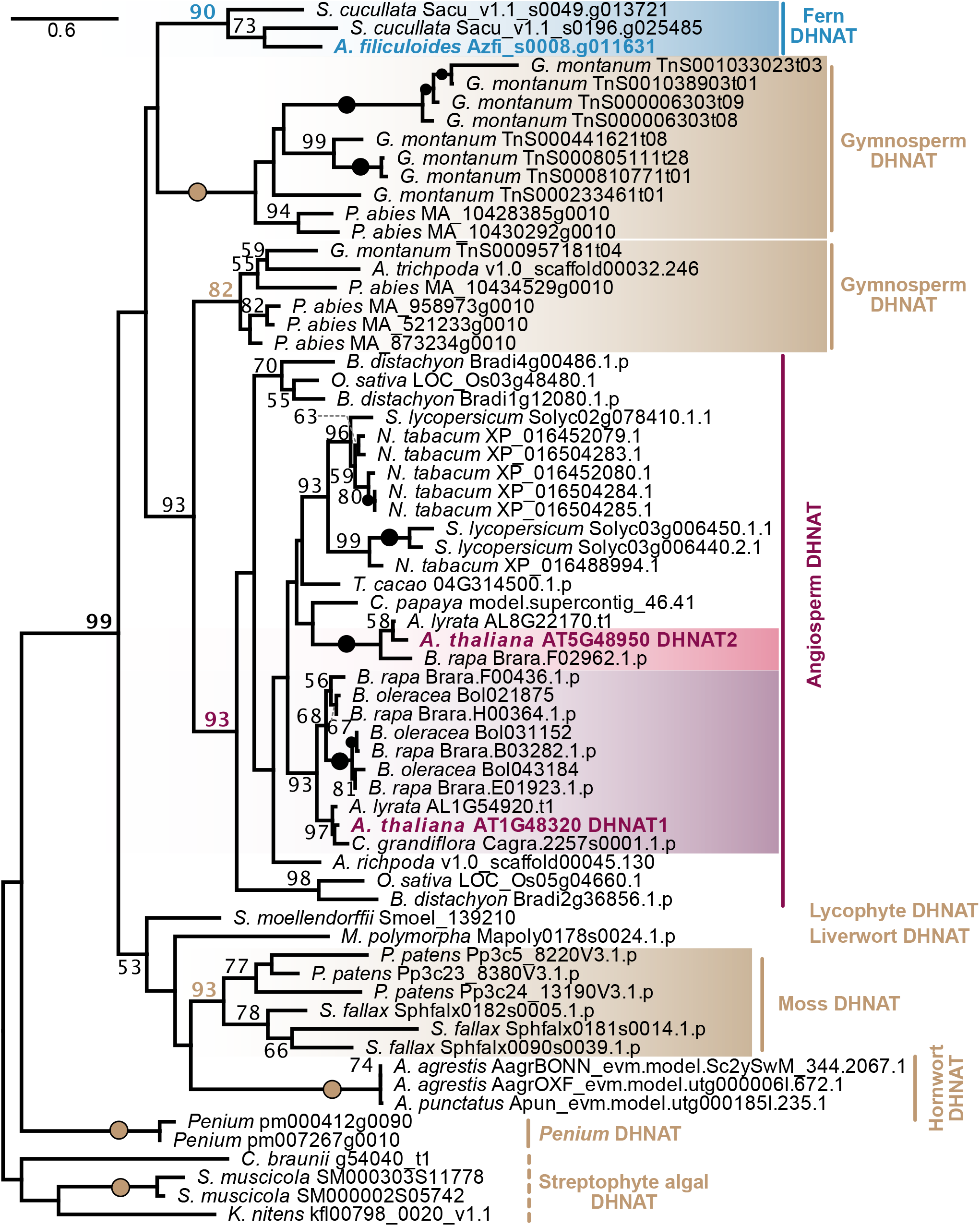
Phylogeny of the DHNAT family. ML-phylogeny was computed with IQ-Tree version multicore version 1.5.5 for Linux 64-bit with 100 bootstrap-replicates; JTT+I+G4 was chosen as model for protein evolution according to Bayesian Information Criterion. Bootstrap support of 100 is indicated by black and colored dots on the individual branches. DHNAT1 and 2 from *A. thaliana* are indicated in purple, the monophyletic DHNAT1 clade is highlighted in purple and the monophyletic DHNAT2 clade is highlighted in pink. DHNAT orthologs in ferns are highlighted in blue and the sequences from *A. filiculoides* is written in blue and bold.

In the non-oxidative pathway, two possible routes towards benzoic acid exist. Both start in the cytoplasm and require a hydratase (which is suggested to occur in both the cytosol and the peroxisomal pathway; and a lyase to synthesize benzaldehyde (Widhalm and Dudareva, 2015). The conversion from benzaldehyde to benzoic acid can occur via two different steps: (i) a cytoplasmic step, catalyzed by AAO4 in *A. thaliana* (Ibdah et al. 2009), or (ii) a mitochondria-localized step, as described for Snapdragon (*Antirrhinum majus*), where BALDH is converting the benzaldehyde to benzoic acid (Long et al. 2009). Given that for the last step towards benzoic acid different routes processed by distinct types of enzymes have been described, we analyzed both of them bioinformatically. Similar to the peroxisomal pathway, BALDH and AAO homologs must have already existed in the common ancestor of all Chloroplastida (Figure 6, S2). While algae contain only one homolog of BALDH, many of the land plant lineages show lineage-specific expansions on BALDH-like homologs (Figure S3). This also includes the fern *A. filiculoides*. In the ALDH2 clade that contains BALDH the fern *S. cucullata* has an ortholog, but not *A. filucloides* (Figure S3). That said, several sequences of *A. filiculoides* fall into the larger clade of ALDH1 and ALDH2 sequences, where they, however, only cluster with weak support (bootstrap value 61) with the ALDH1 rather than ALDH2 sequences. These are therefore the most likely candidates for a BALDH reaction (Azfi_s0021.g015740 [low-confidence sequence], Azfi_s0078.g038198, Azfi_s0049.g030778 and Azfi_s0083.g038897; Figure S3). As for the BALDH-like sequences, the AAO family shows lineage-specific radiation. This is most extensive in angiosperms and the lycophyte *S. moellendorffii* (Figure 6). In contrast, the two ferns have each only one copy for AAO (for *A. filiculoides*: Azfi_s0158.g053886; Figure 6).

**Figure 6.**
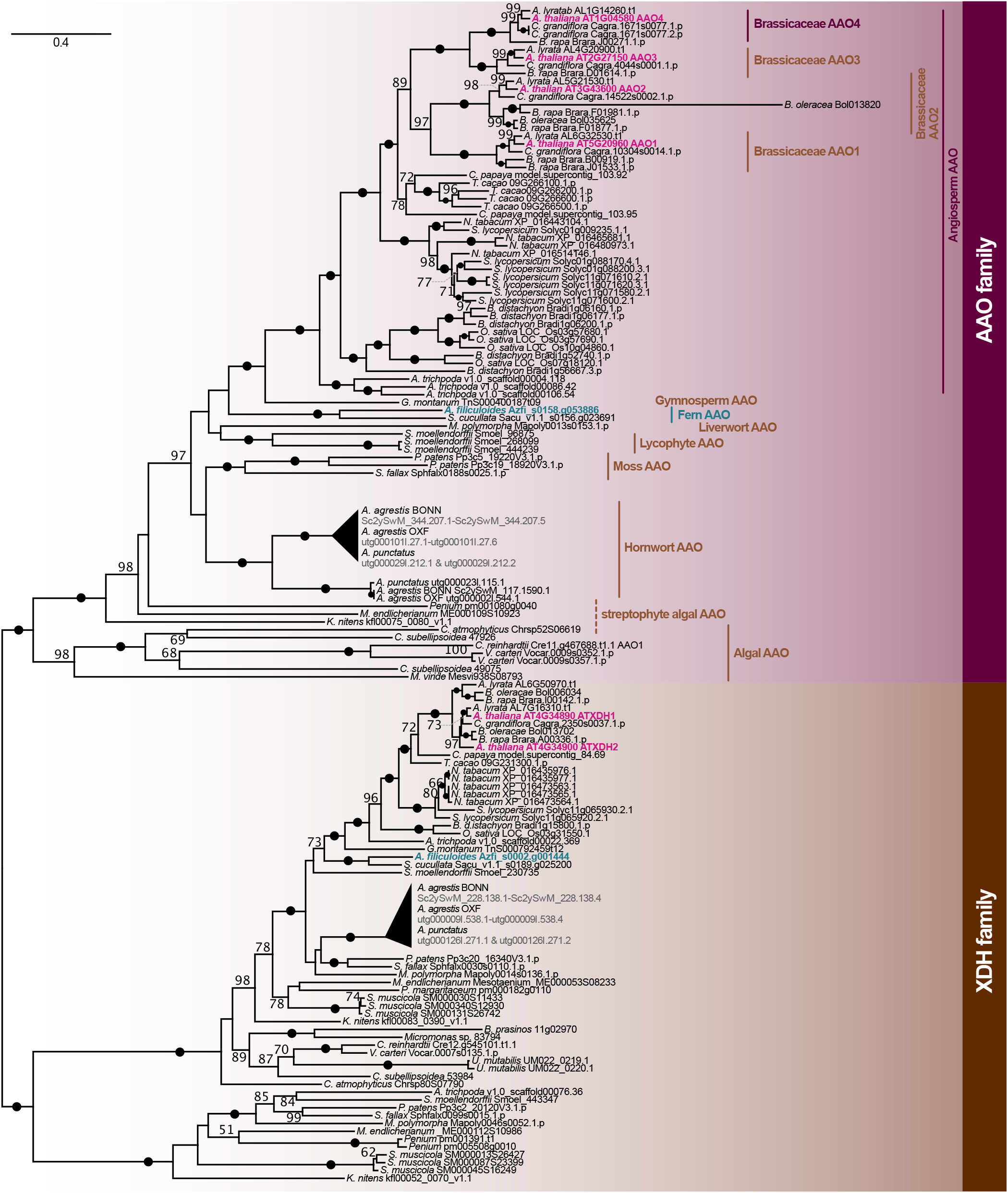
Evolution of the AAO family. ML-phylogeny with 100 bootstrap replicates to recover the evolution of the AAO family; LG+F+I+G4 was chosen as model for protein evolution according to Bayesian Information Criterion. The blastp searches identified sequences either belonging to xanthin dehydrogenase (XDH) and AAO clades. The AAO clade is highlighted in purple and the XDH clade is highlighted in brown. Sequences from *A. thaliana* are indicated in pink and sequences from *A. filiculoides* are indicated in blue. Bootstrap support of 100 is indicated by black dots.

The PAL-dependent pathway is less-explored as a source for SA in land plants compared to ICS-dependent synthesis of SA. The PAL-dependent pathway, despite being highly radiated, has homologs for all enzymes being present at least in the last common ancestor of all streptophytes—going hand in hand with the idea that parts of this pathway were part of the genetic building blocks that allowed for the radiation of spezialized metabolism in land plants (Fürst-Jansen et al., 2020). This is most likely explained by the fact that this pathway is not a highly specialized pathway for SA biosynthesis, but instead recruits steps from other pathways, such as phenylpropanoid. With regard to the water fern *A. filiculoides*, our data adds more support to benzoic acid-derived SA than isochorismate-derived SA biosynthesis (Figure 7). In this unique system, however, another plausible route for synthesizing SA exists. Cyanobacteria from the Nostocaceae, to which the cyanobiont belongs, can synthesize SA (Toribio et al. 2020) and it is possible that *A. filiculoides* has an alternative reservoir for this phytohormone.

**Figure 7.**
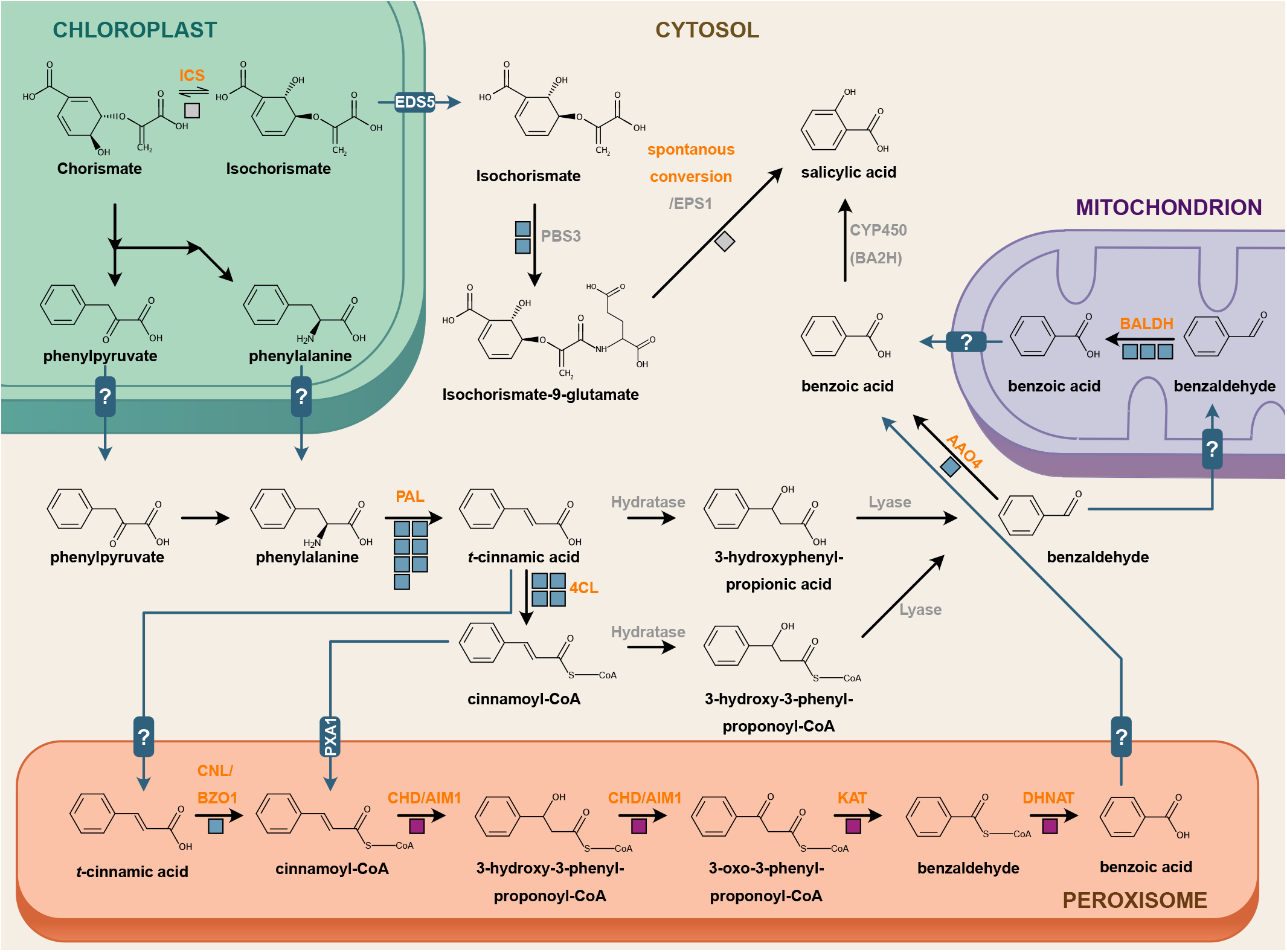
Routes of SA biosynthesis in land plants. SA can be derived from isochorismate or benzoic acid. The route from isochorismate requires ICS and a GH3 enzyme (Wildermuth et al., 2001, Garcion et al. 2008, Rekhter et al. 2019, Torrens-Spence et al. 2019), whereas the benzoic acid-derived SA pathway is less direct requires several steps that function also in other pathways. The reactions are indicated by arrows and the responsible enzymes are written on top of these arrows. Grey enzyme names indicate steps that have not been analyzed in this study because (i) data was available from previous studies or (ii) sequences for these enzymes are not available from functionally characterized enzymes because there is only indirect evidence. Boxes below the arrows indicate the number of high-confidence sequences available in the genome of *A. filiculoides*. Grey boxes indicate absence in *A. filiculoides*, blue boxes indicate the presence of homologs in the fern and pink boxes indicate presence of orthologs encoded in the genome of *A. filiculoides*.

#### The cyanobionts’ pathway

Instead of producing SA autonomously, it is theoretically conceivable that the cyanobiont produces SA *for A. filiculoides*. We therefore investigated the cyanobionts genome for its genetic capability to synthesize SA. Bacteria produce SA either via ICS and IPL enzymes or directly via salicylate synthase (Ankenbauer and Cox, 1988, Gaille et al., 2002, Zwahlen et al., 2007). Indeed, in several Nostocaceae SA has been measured (Toribio et al. 2020). This leaves the possibility that between *Azolla* and its cyanobiont a tight regulation of SA-derived host responses has co-evolved.

We searched for ICS sequences in *T. azollae* using the ICS sequence from *T. variabilis*; we retrieved three sequences annotated as ICS (WP_013190659.1; query coverage 97%, % identity 68.63%, e-value 0), anthranilate synthase component I (WP_013192196.1; query coverage 63%, % identity 27.48%, e-value 2*10^−23^) and aminodeoxychorismate synthase component I (WP_013189914.1; query coverage 71%, % identity 27.54%, e-value 1*10^−28^; Figure 8a). Isochorismate is converted to salicylic acid via IPL (Gaille et al., 2002). Therefore, we next searched for an IPL candidate in *T. azollae*. Using the functionally characterized IPL of *Pseudomonas aeruginosa* (Gaille et al., 2002) as a query for a BLASTp, we found no hits in the genome of *T. azollae* but in several other cyanobacteria, including the order Nostocales and the family Nostocaceae, to which *T. azollae* belongs (Figure 8b). Thus, while some *Nostoc* species are able to synthesize SA (Toribio et al. 2020), only three different species from the Nostocaceae encoded an IPL sequence (Figure 8b).

**Figure 8.**
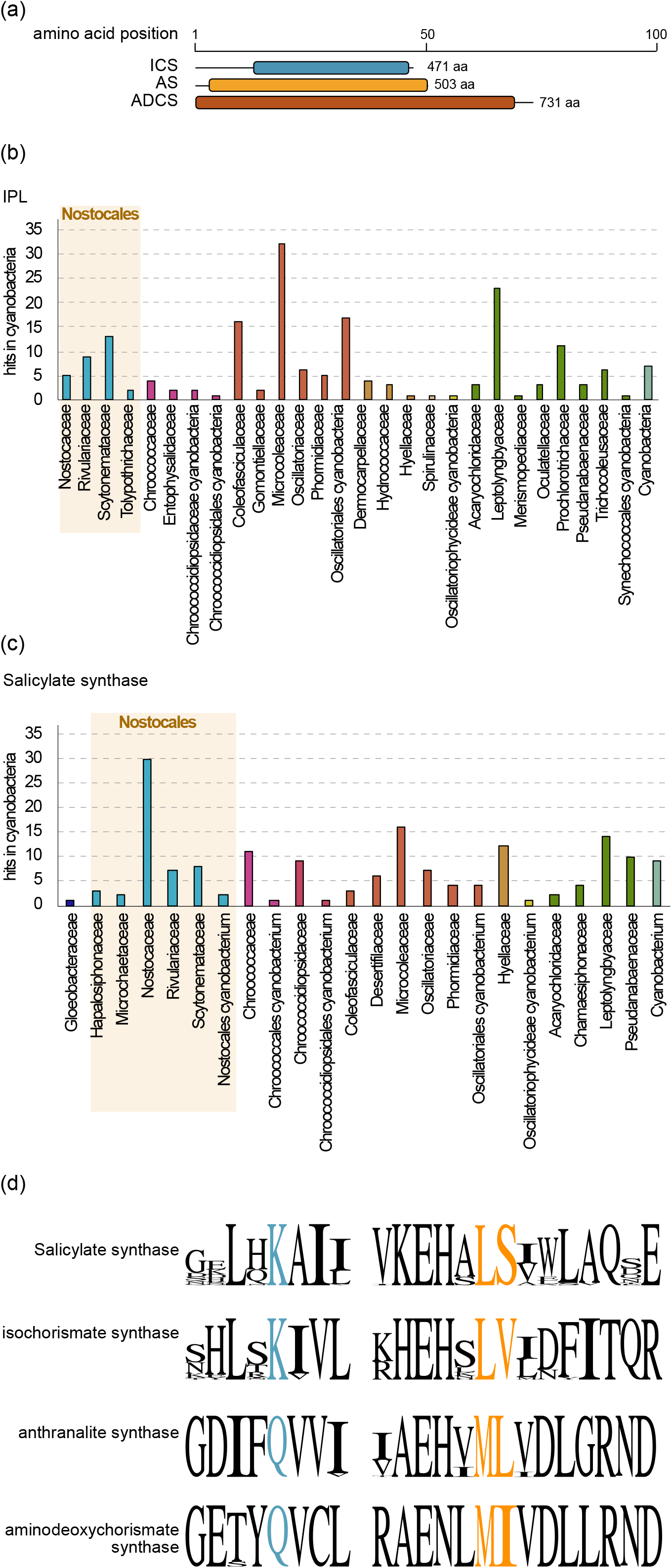
Chorismate-binding enzymes in cyanobacteria. (a) Three chorismate-binding enzymes, an isochorismate (ICS), anthranilate synthase (AS) and aminodeoxychorismate synthase (ADCS), were identified in *T. azollae* using a blastp search. Protein domains identified via TIGR are drawn to scale (blue= isochorismate synthase domain, yellow = anthranilate synthase component I, red = ADCS component I domain). Amino acid positions are indicated by the bar above. (b) Number of blastp hits for IPL in cyanobacteria, using the IPL sequence of *P. aeruginosa* sorted according to taxonomy. Hits recovering sequences from the Nostocales are highlighted by a brown box. (c) Number of hits recovered through a blastp search for salicylate synthase with the sequence from *M. tuberculosis* as query sorted according to cyanobacterial taxonomy. Hits to Nostocales are highlighted by a brown box. (d) Sequence logos for the conserved regions in chorismate binding enzymes. Highlighted in blue and orange are residues relevant for substrate specificity (Plach et al. 2015).

Some bacteria use salicylate synthase, which directly converts chorismate to SA. Therefore, we queried the functionally characterized salicylate synthase from *Mycobacterium tuberculosis* (Zwahlen et al. 2007) in a BLASTp search against (a) *T. azollae* (supplemental data S1A) and (b) all cyanobacteria (supplemental data S1B). In the search against *T. azollae* we retrieved two hits; one against anthranilate synthase component I (e-value 6*10^− 28^), of which the best hit in a protein domain search was indeed against anthranilate synthase component I (TIGR00564, e-value 0) and one against ICS (e-value 2*10^−18^), but no hit to an annotated salicylate synthase. In contrast to the IPL distribution, the most hits of salicylate synthase were in the family of Nostocaceae (Figure 8c). All of the top100 salicylate synthase-like sequence from diverse cyanobacteria, were predicted to contain a salicylate synthase domain (TIGR03494). Yet, all sequences also matched TIGR domain annotations for an ICS domain (TIGR00543), an aminodeoxychorismate synthase component I domain (TIGR00553) and an anthranilate synthase component I domain (TIGR00564). The positions for all domain annotations were partially overlapping. This is in contrast to the two sequences retrieved for *T. azollae*, which had their best TIGR domain hits to an anthranilate synthase component I (TIGR00564, e-value 0) and ICS domain (TIGR00543, e-value 4.75*10^−107^).

As described above engineered anthranilate synthases from *Salmonella* had been described to convert chorismate to isochorismate instead of anthranilate (Plach et al. 2015). Some of these anthranilate synthase mutants even directly synthesized SA from chorismate (Plach et al. 2015). We, thus, had a closer look at the putative chorismate binding enzymes of *T. azollae*. Similar to our analyses for the anthranilate synthase of the host, we aligned the protein sequences of *T. azollae* with that of the anthranilate synthase from *S. enterica* subsp. enterica serovar Typhimurium str. LT2. As described for several major phyla of bacteria (Plach et al., 2015), the anthranilate synthases of cyanobacteria had the QML motif, and the cyanobacterial aminodeoxychorismate synthases encoded the expected QMI motif (Figure 8d). Likewise, salicylate synthase and ICS encoded usual motifs with KLS and KLV, respectively (Figure 8d).

Cyanobacterial anthranilate synthase, including that of *T. azollae*, and aminodeoxychorismate synthase have the QML and QMI motif. Hence, they show the typical aa pattern for these types of enzymes (Plach et al. 2015) and are likely not involved in SA biosynthesis. The ICS had a KLV motif, frequently found for bacterial ICS sequences (Plach et al. 2015). This suggests that *T. azollae* is capable of synthesizing isochorismate but not SA. Given that *Trichormus* species have been shown to produce SA (Toribio et al. 2020), it can be envisioned that *T. azollae* lost its salicylate synthase or IPL. Indeed, during the 66-100 million years of co-evolution between fern and cyanobiont (Hall and Swanson 1968, Collinson 2002; Carrapiço 2006), the genome has started to erode, evident by a high proportion of pseudogenes (Ran et al. 2010). It could therefore, be that due to SA being synthesized by *Azolla, T. azollae* lost its SA synthesizing capabilities. Isochorismate in bacteria can be funneled into the menaquinone pathway (Daruwala et al. 1996, Meganathan and Kwon 2009) or to produce SA under iron deficiency, possibly to act as a siderophore (Visca et al. 1993). In the here described scenario isochorismate of the cyanobiont would singularly be used for menaquinone biosynthesis, while potential SA-associated siderophores may require SA from *Azolla* to be shuffled to its symbiont. However, isochorismate can decompose at slow rates directly into SA and the existence of an IPL may not be necessary when the other isochorismate-metabolizing route towards menaquinone is not active (Rekhter et al. 2019).

#### PAL-dependent routes appear the most likely path towards SA in Azolla

All major land plant lineages and green algae appear to encode an ICS sequence in their genomes (Figure 1a). This includes the only other sequenced fern genome of *Salvinia cucullata—*but not that of *A. filiculoides*. This warrants attention. If we rule out simple technical errors, *A. filiculoides* appears to have secondarily lost its ICS sequence and by that its ability to synthesize isochorismate. It is conceivable that during the evolutionary history of plants and algae, SA biosynthesis had already once been recruited from a cyanobiont—the cyanobacterial plastid progenitor (Gross et al., 2006). In the Brassicaceae *A. thaliana* ICS is localized to the plastid, where it synthesizes isochorismate (Wildermuth et al. 2001, Garcion et al. 2008), which is then transferred to the cytoplasm to be further processed into SA (Rekhter et al. 2019). It is thus tempting to speculate that this is a case where the tape of evolution is replayed: the fern *Azolla* receives isochorismate from its symbiont, *T. azollae*, and uses this backbone molecule to synthesize SA. Indeed, the genome of *T. azollae* shows clear traces of an obligate symbiont and—based on its genome structure and its mode of inheritance—was speculated to be an organelle in the making (Ran et al. 2010). Yet no cases of endosymbiotic gene transfer have been reported for the genome of *A. filiculoides* (Li et al. 2018). Indeed, cyanobiont-free *Azolla* species can survive with ample nitrogen supply (Brouwer et al., 2017). Thus, while it is a fascinating area of speculation, we would suggest that *Azolla* does not have to rely on the cyanobionts for the synthesis of its defense hormone.

Overall, it appears to be more likely that *Azolla* uses the PAL-dependent pathway. This does not exclude that *T. azollae* requires host-derived SA. Indeed, that both partners rely on the phytohormone to some degree is in agreement with our previous data, where exogenous application of an SA-derivative triggered alterations in host and cyanobiont gene expression patterns as well as cyanobiont abundance (de Vries et al. 2018). The obvious question following this is, whether SA biosynthesis and SA response can be modulated by the cyanobionts. To gain first insights, we next investigated the expression profile and differential gene expression (DGE) of the PAL-dependent biosynthesis pathway and SA responses of *A. filiculoides* upon changes in nitrogen availability and presence/absence of the cyanobiont.

### Alterations in the cyanobionts population influence the gene expression patterns for benzoic acid-derived SA biosynthesis

Nitrogen availability has been shown to modulate expression of genes involved in the synthesis of phenylpropanoid-derived flavonoids in *A. filiculoides* (Güngör et al. 2021). Indeed, the absence of cyanobionts also change expression levels of *CHS* in *A. filiculoides* (Eily et al. 2019), overall linking the phenylpropanoid-derived compounds to the symbiosis. SA, which is a conceivable phenylpropanoid-derived compound in *A. filiculoides*, is also a regulator of flavonoids (Nugroho et al. 2002, Tounekti et al. 2013, Ni et al. 2018). In bacteria, SA is likely used as a siderophore to sequester iron from iron-poor environments (Visca et al. 1993). Furthermore, in 2018 we have shown that SA can influence the population size of the cyanobionts and their nitrogen fixation-related gene expression (de Vries et al., 2018). To explore these factors, we (a) investigated the gene expression profiles upon these two factors and (b) calculated the DGE for *A. filiculoides* with resepct to the PAL-dependent pathway.

To understand the global gene expression patterns of genes for SA synthesis and down-stream pathways in *A. filiculoides*, we first analyzed gene expression levels in Transcript Per Million (TPM, supplemental data S2A and S3A). In total, we investigated 19 high-confidence candidate genes; 18 of these showed an average TPM above 1 in at least one condition. *A. filiculoides* invested a pronounced transcript budget into the phenylpropanoid pathway: while the cumulated expression of all genes amounted to an average (over all conditions) of 2734.79±544.91 TPM, the 11 homologs of genes from the canonical phenylpropanoid pathway (7 *PAL-*, and 4 *4CL*-homologs) made up 2335.55±535.27 TPM—which was largely due to the PAL homologs. Of the seven putative PAL-coding genes, one distinguished itself (*Azfi_s0123*.*g048412*) by dwarfing all the other average gene expression levels with a TPM of 1757.05±338.51. At the same time, *Azfi_s0123*.*g048412* had the most stable expression level, never showing differential gene expression changes under the conditions analyzed here. For comparison, the TPM of 4CL amounted to 24.44±11.58.

11 of 19 high-confidence candidate genes for the SA biosynthesis pathway in *A. filiculoides* showed significant changes in their differential gene expression patterns due to supply of fixed nitrogen and/or presence of the cyanobiont (Benjamini Hochberg adjusted FDR ≤ 0.05; Figure 9a based on the pathways shown in Figure 7, Supplemental data S2B); 8 of these 11 genes were up or down-regulated with a log_2_(fold-change[FC]) > 1 or <-1. The absence of fixed nitrogen alone in presence of the cyanobiont (nYcY vs. nNcY) or the absence of the cyanobiont in presence of fixed nitrogen (nYcY vs. nYcN) has only a small effect on the expression of the PAL-dependent biosynthesis pathway: only 3 of 19 genes showed differential gene expression in either of the comparisons. Most changes were observed when comparing the presence of cyanobionts (nNcY) to the absence of cyanobionts (nNcN) in the absence of fixed nitrogen (8/19 differentially expressed genes) and when comparing presence of fixed nitrogen and cyanobiont (nYcY) with the absence of both (nNcN; 9/19 differentially expressed genes). These two comparisons shared seven differentially expressed genes, of which two (belonging to gene families *PAL* and *BALDH*) were up and five were down-regulated (belonging to gene families *PAL, 4CL, BALDH* and *AAO4*) in both conditions.

**Figure 9.**
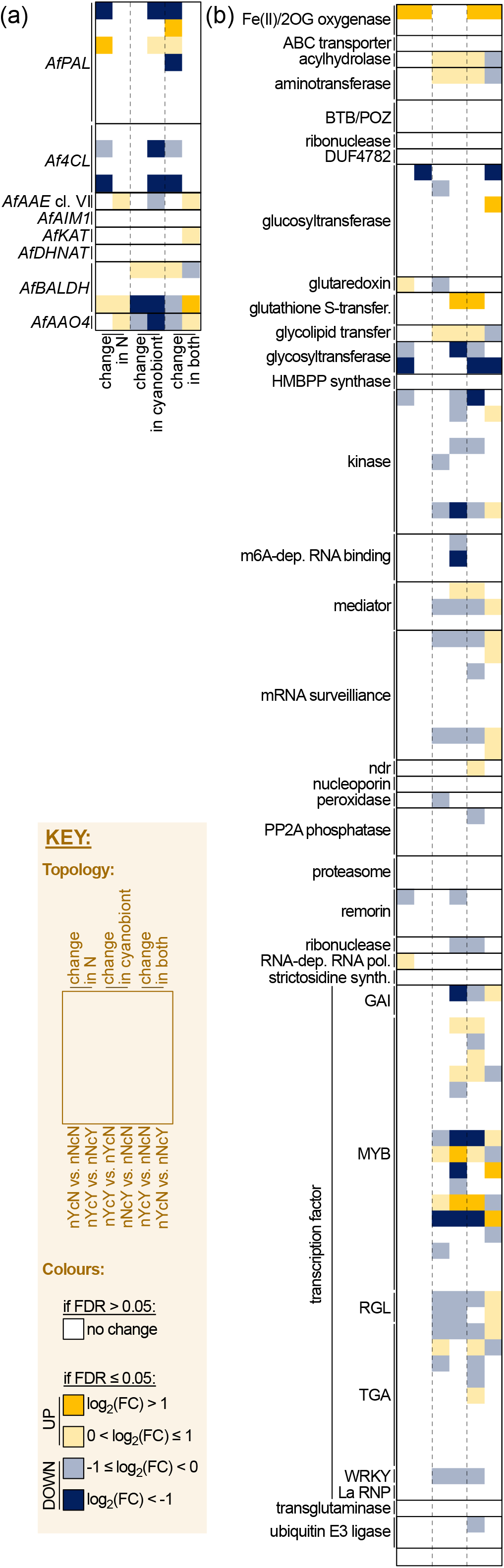
Differential gene expression of homologs of genes for SA biosynthesis and SA responses. (a) Differential gene expression of genes associated with the PAL-dependent pathway for SA biosynthesis in *A. filiculoides*, including candidates for *AfPAL, Af4CL, AfAAE clade VI* (i.e. the closest homologs to *AtBOZ1* and *PhCNL* in *A. filiculoides*), *AfAIM1, AfKAT, AfDHNAT, AfBALDH*, and *AfAAO4*. (b) Differential gene expression of homologs of SA-responsive genes. Significant differential regulation is indicated by blue and yellow colors. Light yellow indicates significant differential up-regulation (FDR ≤ 0.05) with a log_2_(FC) above 0 and below 1, dark yellow indicates significant differential up-regulation (FDR ≤ 0.05) with a log_2_(FC) above 1. Light blue indicates differential significant down-regulation (FDR ≤ 0.05) with a log_2_(FC) below 0 but above −1 and dark blue indicates significant down-regulation (FDR ≤ 0.05) with a log_2_(FC) below −1.

The two initial steps in the core phenylpropanoid pathway, *PAL* and *4CL*, the initial step in the β-oxidative pathway (*BZO1*) and the last steps in the conversion of cytoplasmic benzaldehyde to benzoic acid (*AAO4* and *BALDH*) were responsive on a gene regulatory level (Figure 9a). The peroxisomal steps catalyzed by gene families *AIM1, DHNAT1/2* and *KAT* showed little differential gene expression; only the *AfKAT* candidate was minimally induced (log_2_(FC)=0.38; FDR=0.02) when both nitrogen and cyanobiont conditions were changed (nYcN vs. nNcY). This is in agreement with the phenylpropanoid pathway being regulated on a transcriptional level (e.g. Weisshaar and Jenkins 1998, Deluc et al. 2006, Xie et al. 2008). *PAL* genes are responsive to biotic stressors (Oliva et al. 2015, Reboledo et al. 2021). It is therefore, in general, not surprising that genes from the phenylpropanoid pathway are responsive to the treatment. In addition, the late steps in the pathway are modulated due to the availability of the cyanobiont and fixed nitrogen. Hence not only SA appears to alter the abundance of the cyanobiont and expression of genes relevant for nitrogen fixation (de Vries et al. 2018), availability of both components also influences SA biosynthesis. This suggests an intricate system of communication between the two partners.

The pattern of how SA biosynthesis is regulated on the gene level appears complex. Sometimes it is contrasting within a gene family or opposite responses occur when changes in cyanobiont presence/absence is evaluated compared to when changes in the availability of fixed nitrogen are evaluated. It appears that in the presence of cyanobionts more genes are down-regulated than up-regulated, though (Figure 9a). Assuming that down-regulation of genes in the SA biosynthesis pathway translates into less SA being produced, this appears to be in contrast with SA levels being induced during colonization of some symbionts in other plants (Fernández et al. 2014). However, it was noted that such defenses are early on responses, hypothesized to stop symbionts from too much proliferation (Plett and Martin 2018). An integrated symbiont, however, is a long-term association. Thus, using SA to limit proliferation within a host early on during symbiosis is not relevant. Moreover, in our previous data on exogenous application of MeSA, we observed an increase of the cyanobiont after treatment (de Vries et al. 2018). Taken together with the expression data on the putative SA biosynthesis genes one may hypothesize a feedback loop between the accumulation of SA and the accumulation of the cyanobiont. This is in agreement with MeSA reducing the expression of the gene encoding iron-dependent niFE (de Vries et al. 2018).

### Expression profile of SA responsive pathways in *A. filiculoides* show distinct responses towards the availability of fixed nitrogen and the cyanobiont

In several land plant species, SA levels are induced during interactions with beneficial microbes (Blilou et al. 1999, Blilou et al. 2000, Liu et al. 2003, Pozo et al. 2015). Further cyanobacterial treatment of *A. thaliana* resulted in the induction of SA controlled gene expression (Belton et al. 2020). Above, we showed that gene expression levels associated with SA biosynthesis are reduced in presence of the cyanobionts. Hence, we wondered whether SA responsive genes may also differ in their gene expression in this particular symbiosis compared to less intimate ones.

We downloaded all *A. thaliana* protein sequences of genes that are associated with the term ‘SA responses’ from TAIR (Lamesch et al., 2011). These 176 sequences were used as a query in a reciprocal BLASTp approach, first against all proteins of *A. filiculoides* and the thusly obtained homologs back against *A. thaliana*. In total we found 105 homologs, of which 97 are high confidence proteins. These 97 homologs correspond to 47 different SA-responsive proteins of *A. thaliana* involved in various processes (Figure 9b, Supplemental data S3B). Overall, 55.7% of the homologs of SA-responsive genes show differential gene expression (FDR<0.05) in at least one comparison. Similar to the SA biosynthesis expression, SA-derived responses showed most differential expressed genes in the comparisons nNcY vs. nNcN (31/97 homologs) and nYcY vs. nNcN (35/97 homologs). Of those genes, 23 were shared between the conditions, nine were up-regulated and 14 down-regulated (FDR<0.05). Corroborating the results of the expression patterns of the SA biosynthesis, changes in the availability of fixed nitrogen have only little influence on the expression of SA responsive genes. Independent of the presence of the cyanobiont only two (nYcY vs. nNcY) and seven genes (nYcN vs. nNcN) were significantly changed in their expression (FDR <0.05). Of those only one (*Azfi_s0427*.*g069291*) showed the same direction of regulation, and was induced in both treatments (log_2_-foldchange=1.64 in nYcY vs. nNcY and log_2_-foldchange=2.31 in nYcN vs. nNcN). *Azfi_s0427*.*g069291* was also induced in comparisons where presence of fixed nitrogen and cyanobiont was changed (FDR<0.05). *Azfi_s0427*.*g069291* encodes a putative 2-oxoglutarate (2OG)/Fe(II)-dependent oxygenase (2-ODD) superfamily protein. Given the diversity of reactions in specialized metabolism and hormone homeodynamics that are catalyzed by 2-ODDs it is difficult to make an educated prediction of what the regulation of this 2-ODD means. That said, it is an interesting candidate enzyme for future studies.

Overall, presence of nitrogen compared to its absence and presence of the cyanobiont compared to its absence result in more genes associated with a response to SA being down-regulated than up-regulated (Figure 9b). This indicates that SA signaling is reduced in the presence of fixed nitrogen. The observation was more pronounced when the presence of the cyanobiont was altered than when the availability of fixed nitrogen was changed. This is due to the generally low transcriptional response to changes in fixed nitrogen. In contrast to the abundance of down-regulated genes in response to a change of one component in the system, changes in both components lead to a complex dysregulation of putative SA-responsive genes (Figure 9b).

Many of the differentially expressed genes are found in the category of transcription factors and MAP kinases. Here most homologs appear down-regulated when the presence of the cyanobiont is compared to its absence, independent of the availability of nitrogen (FDR <0.05, Figure 9b). In the comparison of nYcY vs. nYcN eight *TFs* are down-regulated and only three are up-regulated. Also, two *MAP kinase* homologs are down-regulated and none are up-regulated. In nNcY vs. nNcN 10 *TFs* and four *MAP kinases* show down-regulation, while five *TFs* and no *MAP kinase* show up-regulation. It appears that especially up-stream regulators of defense responses are affected by the presence of the cyanobiont. This is in agreement with the reduction of gene expression of the putative SA biosynthesis pathway we observed in the same comparisons (Figure 9a). In contrast, SA signaling was induced upon treatment of *A. thaliana* with *Nostoc punctiforme* PCC 73102 (Belton et al. 2020), a cyanobacterium from the same family as *Azolla*’s cyanobiont. *N. punctiforme* PCC 73102 is a strain with symbiotic competence that is often used in experiments with cyanobacteria and different hosts (Meeks et al. 2001, Warshan et al. 2017). This fits the hypothesis of Plett and Martin (2018), that early increase in SA during symbioses may be to establish the borders of same associations. However, the association between *T. azollae* and the water fern is perpetual. It is thus to be expected that the usual early responses to encountering potential symbionts are not mirrored in this vertically inherited association. Besides, data on host-symbiont interaction is often gathered from angiosperm systems. Data related to SA signaling in other plants during interaction with pathogenic microorganisms already indicate differences in the defense network; for example, the antagonism between SA and another phytohormone, jasmonic acid, appears to not exist in spruce during infection with a necrotrophic pathogen (Arnerup et al. 2013). Additionally, it appears that the genes coding for the Arabidopsis SA receptors *At*NPR1, 3 and 4, which have contrasting roles in SA signaling (Ding et al. 2018), arose from a lineage-specific radiation (Li et al. 2020, supplemental data therein). In *A. filiculoides* two possible candidates for *NPR* homologs, likely derived from a lineage-specific duplication, are found (Li et al. 2020, supplemental data therein). Euphyllophytes thus likely had only a single ancestral NPR protein. Given these data, it is reasonable to assume that SA signaling is in *A. filiculoides* in general and in the symbiosis of the fern only partially similar to that of model angiosperms.

## Conclusion

The phytohormone SA is important for biotic interactions in plants and iron-sequestration in bacteria. Previously, we showed that SA is a candidate molecule involved in the tight coordination of the symbiosis between *A. filiculoides* and its cyanobionts (de Vries et al., 2018). Here, we investigated the possible routes by which SA may be synthesized in *A. filiculoides* as well as how the cyanobiont and the availability of fixed nitrogen influence gene expression patterns that can be associated with SA biosynthesis and signaling. Our data pinpoint that the most likely biosynthetic route towards SA in the fern derives from benzoic acid, rather than isochorismate. Although the cyanobiont is genetically capable of synthesizing the latter, it lacks genes encoding IPL or salicylate synthase to convert isochorismate or chorismate to SA. Most genes relevant for SA biosynthesis in the water fern as well as homologs to genes associated with response to SA are down-regulated in *A. filiculoides* with a cyanobiont compared to a *Trichormus*-free culture. Most of the differential regulation appears to be on up-stream regulators such as *TFs* and *MAP kinases*. Taken our previous and new data together, SA might participate in a feedback loop between *A. filiculoides* and *T. azollae*.

## ACKNOWLEDGEMENTS

JdV thanks the European Research Council for funding under the European Union’s Horizon 2020 research and innovation programme (Grant Agreement No. 852725; ERC-StG “TerreStriAL”). IF gratefully acknowledges support by the Deutsche Forschungsgemeinschaft (INST 186/822-1).

