## Supplemental Figure S1 for "An ancient route towards salicylic acid and its implications for the perpetual *Trichormus–Azolla* symbiosis"

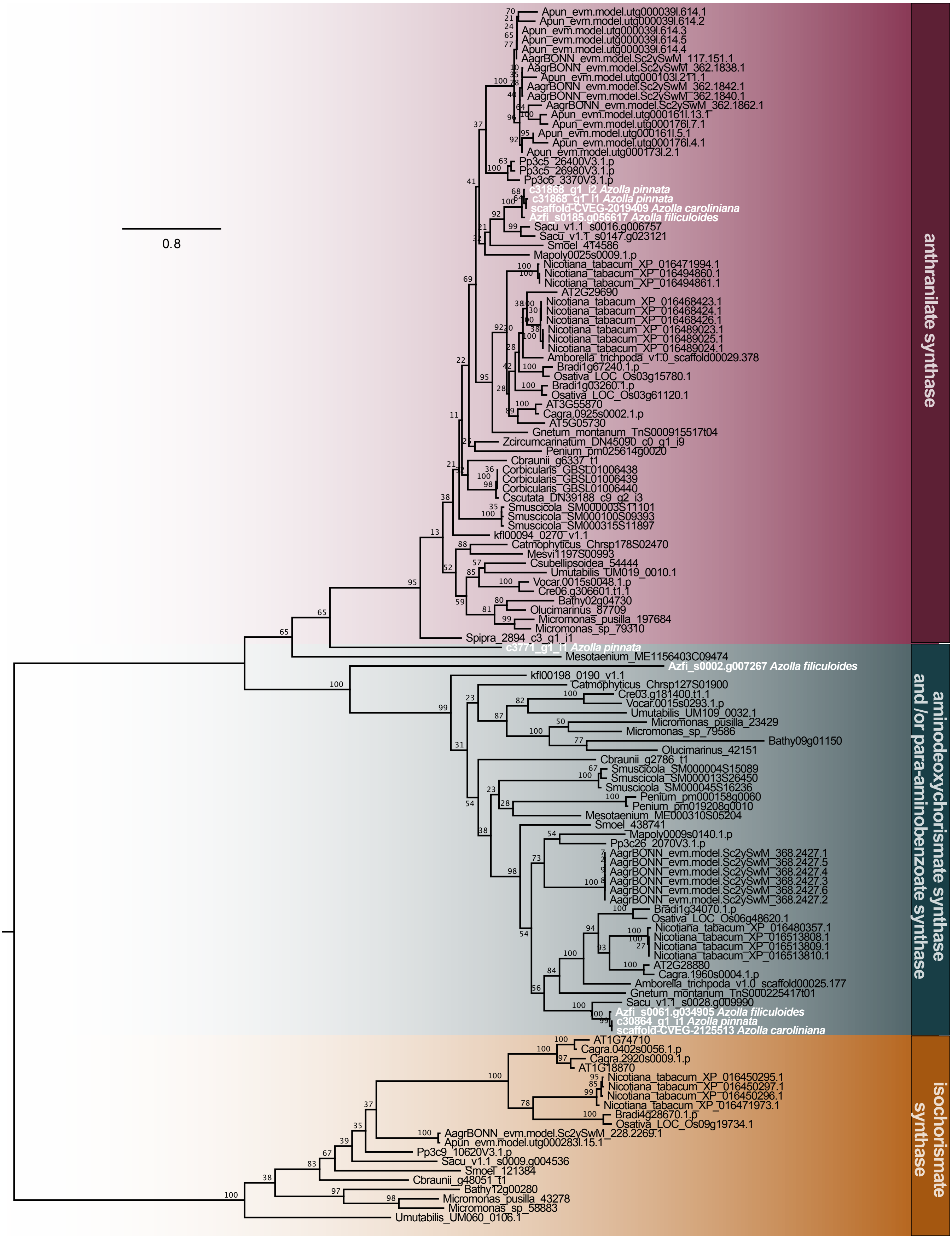


**Figure S1. Phylogeny of chorismate binding enzymes including other *Azolla* spp.** ML-phylogeny of chorismate binding enzymes from Chloroplastida with 100 bootstrap replicates. We used ModelFinder to determine the best model for protein evolution and picked LG+F+I+G4 according to Bayesian Information Criterion. Sequences from *A. thaliana* are highlighted in pink and sequences of *A. filiculoides* are highlighted in blue. We recovered three clear clades, one for isochorismate synthases (ICS), one for aminodeoxychorismate synthases and *p*-aminobenzoate synthases and one for anthranilate synthases.
