## Supplemental Figure S2 for "An ancient route towards salicylic acid and its implications for the perpetual *Trichormus–Azolla* symbiosis"

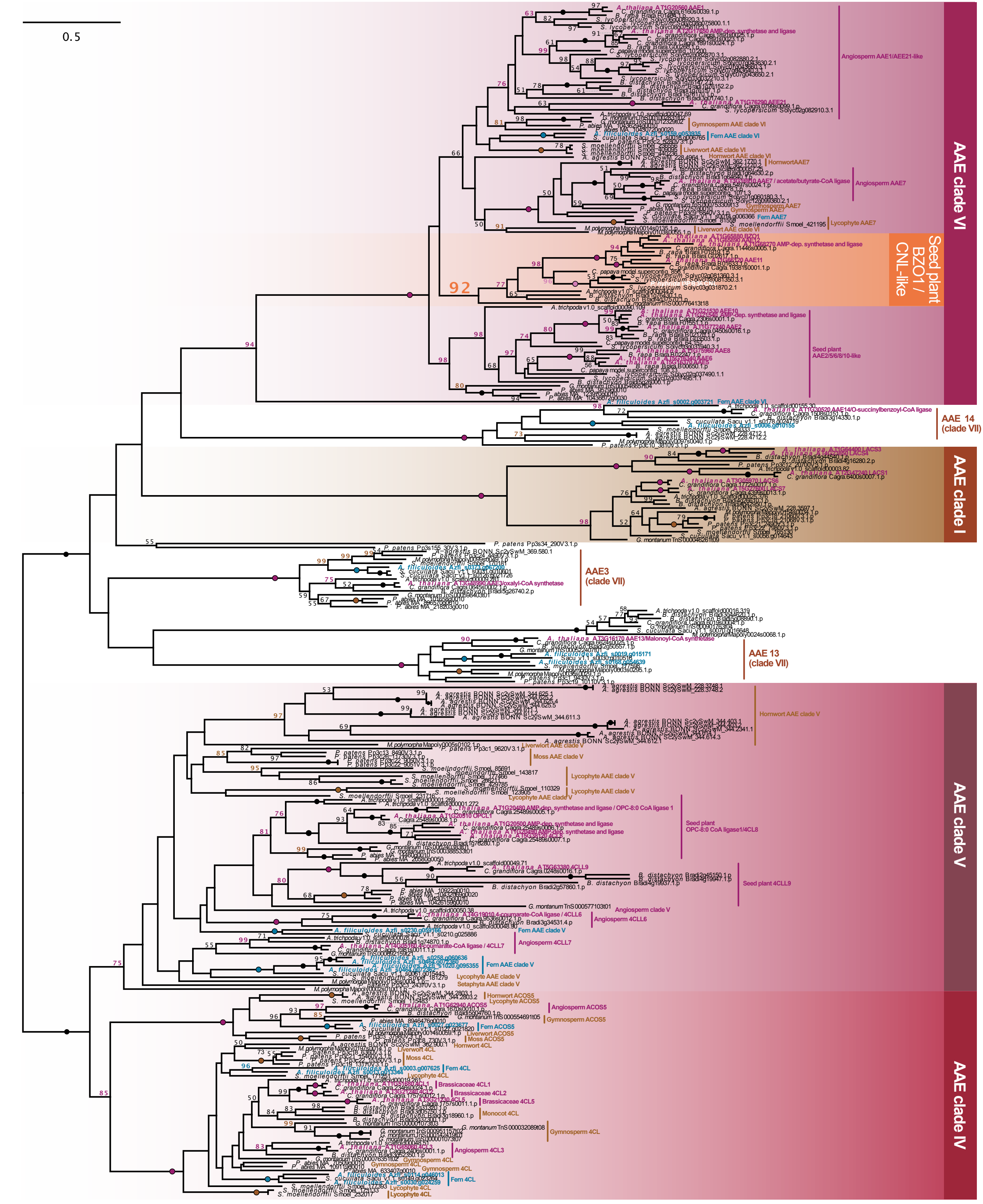


**Figure S2. Phylogeny of Acyl Activating Enzymes, including BZO1.** ML-phylogeny of Acyl Activating Enzymes (AAE) from Chloroplastida with 100 bootstrap replicates. We used ModelFinder to determine the best model for protein evolution and picked LG+I+G4 according to Bayesian Information Criterion. Sequences from *A. thaliana* are highlighted in pink, sequences of *A. filiculoides* are highlighted in blue, and *Petunia hybrida* sequences are highlighted in white font. The BZO1 clade, which is nested within AAE clade IV, is highlighted in orange.
