## Supplemental Figure S3 for "An ancient route towards salicylic acid and its implications for the perpetual *Trichormus–Azolla* symbiosis"

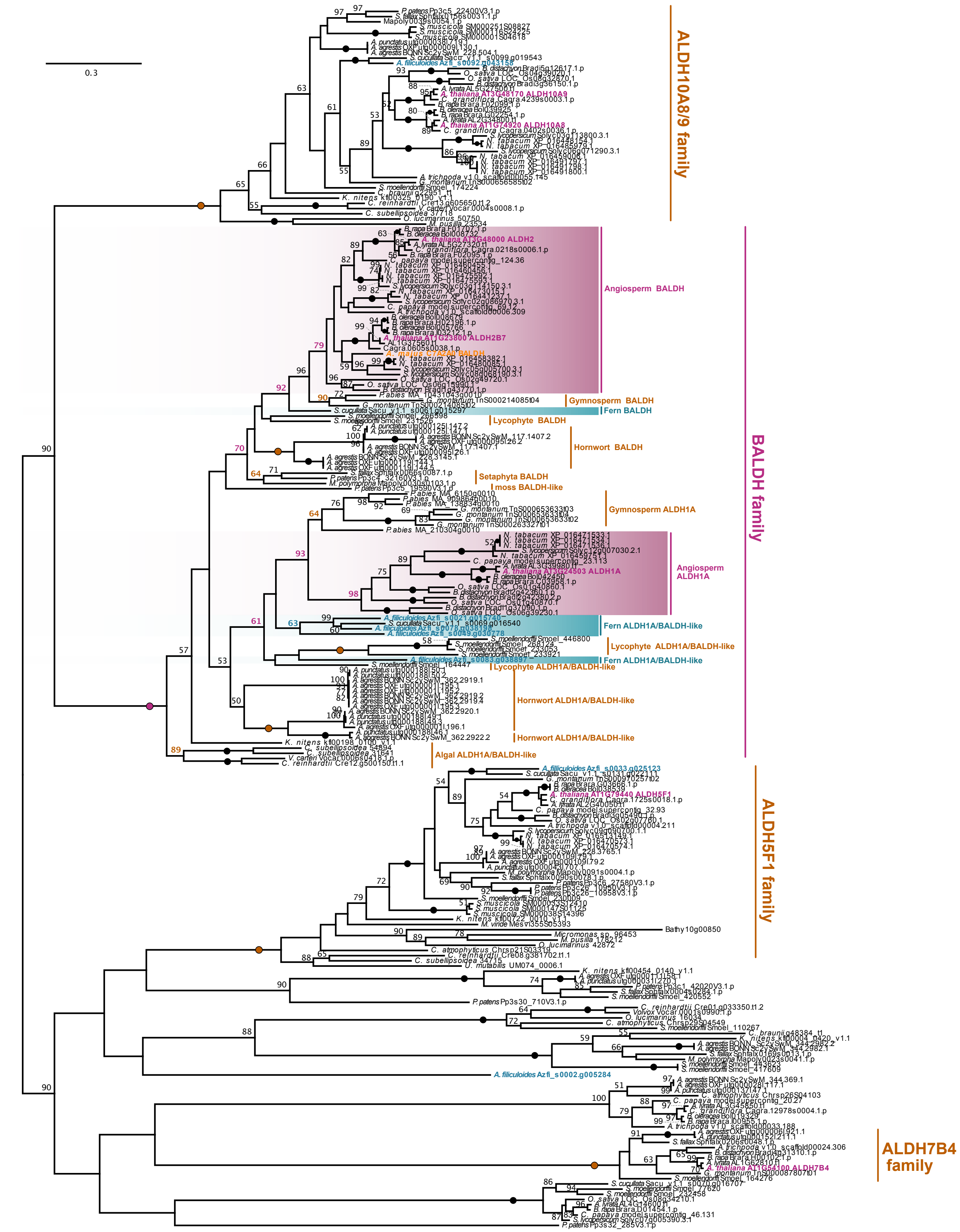


**Figure S3. Phylogeny of Aldehyde Dehydrogenase (ALDH), including Benzaldehyde Dehydrogenase (BALDH).** ML-phylogeny of ALDH from Chloroplastida with 100 bootstrap replicates. We used ModelFinder to determine the best model for protein evolution and picked LG+I+G4 according to Bayesian Information Criterion. Sequences from *A. thaliana* are highlighted in pink, sequences of *A. filiculoides* are highlighted in blue, and the Snapdragon BALDH sequence is highlighted in orange font.
